# Boosting variant-calling performance with multi-platform sequencing data using Clair3-MP

**DOI:** 10.1101/2023.05.31.543184

**Authors:** Huijing Yu, Zhenxian Zheng, Junhao Su, Tak-Wah Lam, Ruibang Luo

## Abstract

**Background:** With the continuous advances in third-generation sequencing technology and the increasing affordability of next-generation sequencing technology, sequencing data from different sequencing technology platforms is becoming more common. While numerous benchmarking studies have been conducted to compare variant-calling performance across different platforms and approaches, little attention has been paid to the potential of leveraging the strengths of different platforms to optimize overall performance, especially integrating Oxford Nanopore and Illumina sequencing data.

**Results:** We investigated the impact of multi-platform data on the performance of variant calling through carefully designed experiments with a deep learning-based variant caller named Clair3-MP (Multi-Platform). Through our research, we not only demonstrated the capability of ONT-Illumina data for improved variant calling, but also identified the optimal scenarios for utilizing ONT-Illumina data. In addition, we revealed that the improvement in variant calling using ONT-Illumina data comes from an improvement in difficult genomic regions, such as the large low-complexity regions and segmental and collapse duplication regions. Moreover, Clair3-MP can incorporate reference genome stratification information to achieve a small but measurable improvement in variant calling. Clair3-MP is accessible as an open-source project at: https://github.com/HKU-BAL/Clair3-MP.

**Conclusions:** These insights have important implications for researchers and practitioners alike, providing valuable guidance for improving the reliability and efficiency of genomic analysis in diverse applications.

## Background

The increasing affordability of sequencing technologies and the rapid evolution of variant-calling methods has opened the door to more research and applications for small variant calling in the human genome (1). It has become more common to possess sequencing data from different sequencing platforms to address a broader range of biological problems (2). Nowadays, the general approach to sequence a genome is through next-generation sequencing (NGS) technology and third-generation sequencing (TGS) technology. The distinct error profiles and read length on different sequencing platforms have prompted benchmarking and discussion of variant calling using different sequencing data (3). Reads generated from NGS technology, like Illumina, are usually short (about 250 bp) and accurate. The methods developed for variant calling using Illumina data, such as GATK (4), have shown promising results in most of the human genome regions. However, due to the read length limitation, in regions that contain duplications longer than 1k bp, it is difficult to further improve variant calling with NGS reads, even with the development of new methods (3). On the other hand, TGS technology like Oxford Nanopore Technologies (ONT) and Pacific Biosciences (PacBio) generate reads ranging from thousands to hundreds of thousands of bases. TGS can produce longer sequences across repetitive and complex genomic regions to which Illumina reads have difficulty being mapped accurately (5), providing better coverage for the genome. DeepVariant (6) and Clair3 (7) are two leading deep-learning-based methods developed for variant calling supporting TGS data. However, ONT data provides lower indel-calling performance due to lower read accuracy, especially in the regions containing homopolymers (3). Therefore, researchers often utilize different platform data for different needs (3).

There are studies that leverage the strength of different platforms to maximize the performance of variant calling (8). The DeepVariant hybrid model is the first variant caller that reported a case study for combining PacBio and Illumina sequencing data (6). It has shown improvements compared to variant-calling performance using single-platform sequencing data (6). Ratatosk, a later published method for error correction, corrects ONT long reads with short reads using a de Bruijn graph to carry information between the two technologies, thus leveraging the benefits of both (9). However, to the best of our knowledge, there is no thorough study investigating the integration of ONT-Illumina data and ONT-PacBio data in a deep-learning model for variant calling. On the other hand, although benchmarking for variant-calling approaches has revealed that ONT, Illumina PacBio data can lead to various types of performance in different genomic regions or at various coverage, the performance in these circumstances when data from different error profiles and read lengths are combined remains unanswered.

Here, we present Clair3-MP (Multi-Platform) (Fig 1), which processes multi-platform data combinations, including ONT-Illumina, ONT-PacBio, and PacBio-Illumina, to perform variant calling. Clair3-MP features a neural network that supports multi-platform data and trains a series of new models, tailored to perform variant calling using different multi-platform data. In addition, Clair3-MP can incorporate reference genome stratification information by including a stratification channel in its input tensors. This channel encodes the platform preference into the neural network and enables better variant-calling performance for multi-platform data.

**Fig. 1.**
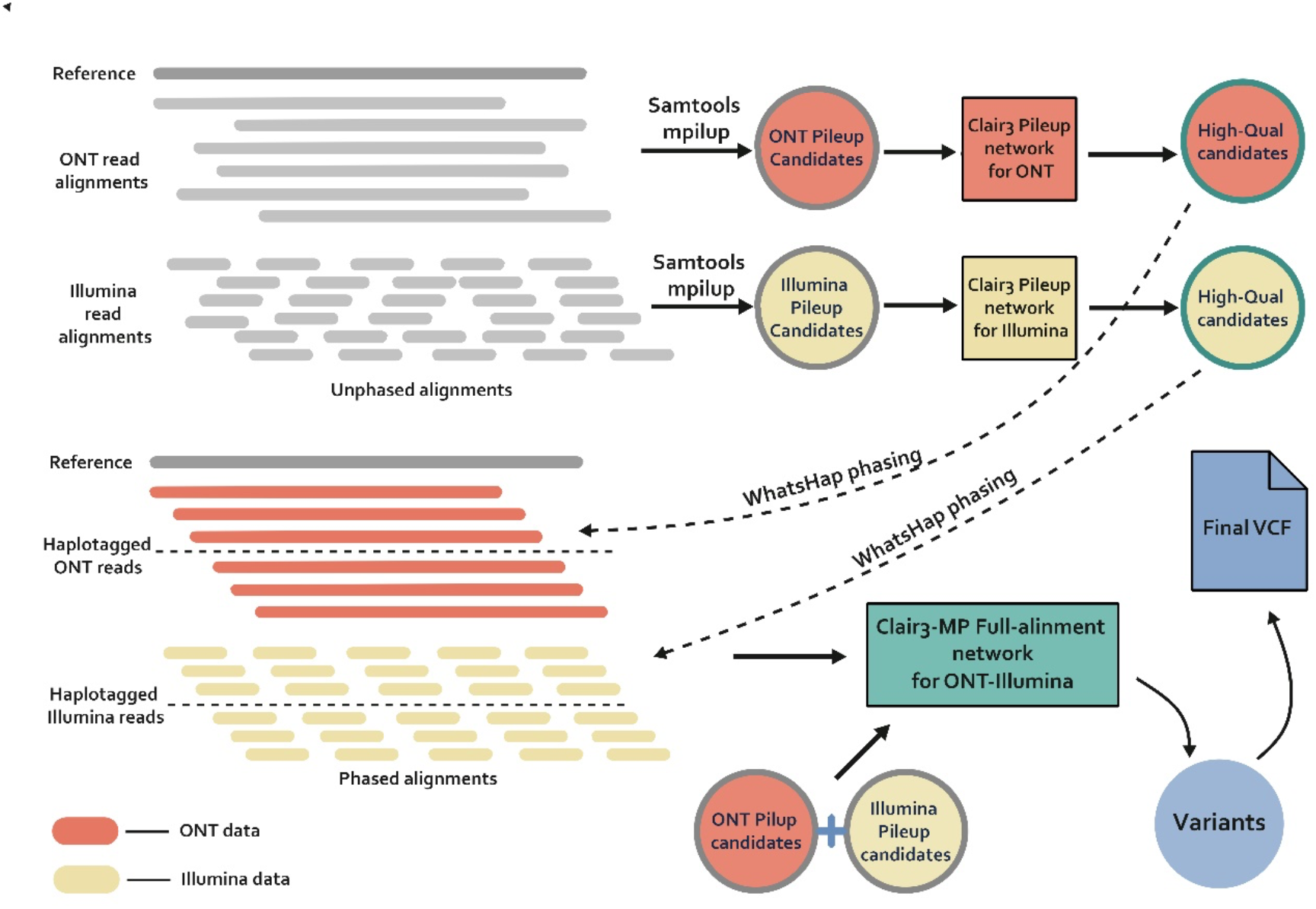
Workflow of Clair3-MP with the ONT-Illumina module. We use ONT-Illumina as an example for calling using Clair3-MP. The platform mentioned in this example can be used on other platforms (e.g., Illumina, PacBio, and ONT). First, candidates from Samtools mpileup are extracted for both ONT and Illumina. Then, the ONT and Illumina pileup candidates are processed independently by the Clair3 pileup models using its respective and recommended parameters. The resulting high-quality candidates are used to phase the alignments in its respective platform, using WhatsHap. Finally, all the pileup candidates from the first step and the phased alignments are processed by the Clair3-MP model trained for ONT-Illumina data.

Through our experiments integrating ONT and Illumina reads in Clair3-MP, we observed that sufficient coverage of ONT data (i.e., 30x) improves variant-calling performance compared to using only data from either platform. Variant calling using multi-platform data containing 30x ONT and Illumina data at 10x, 20x, and 30x can achieve an F1 score of +0.0545, +0.0058, and +0.0010, respectively, compared to Clair3 with Illumina data at 10x, 20x, and 30x. However, the improvement is not consistent when there is less ONT data in the multi-platform data. Therefore, we recommend performing variant calling using Clair3-MP with ONT-Illumina data with at least 30x coverage of ONT data. On the other hand, variant calling using multi-platform data containing 10x Illumina and ONT data at 10x, 20x, and 30x can achieve an F1 score of +0.0379, +0.0520, +0.0545, respectively, compared to Clair3 with 10x Illumina data. This suggests that with low coverage of Illumina data, additional ONT data at any coverage can guarantee improved variant calling with Clair3-MP. We further stratified the variant-calling results of Clair3-MP with 30x ONT and 30x Illumina and compared them to Clair3 with 30x ONT or 30x Illumina to observe the area of improvement in a setting that contains enough data for both platforms. The stratification analysis reveals that the increased variant-calling performance in Clair3-MP with ONT and Illumina data stems from an improvement in difficult genomic regions. We observed that there are increased SNP and Indel F1 scores of Clair3-MP with ONT and Illumina data compared to Clair3 with ONT or Illumina in these difficult regions: large low-complexity regions (SNPs: from 0.9963 or 0.9844 to 0.9973, Indels: from 0.9392 or 0.9661 to 0.9679), segmental duplication regions (SNPs: from 0.9565 or 0.9177 to 0.9653, Indels: from 0.9022 or 0.9300 to 0.9566), and collapse duplication regions (SNPs: from 0.7797 or 0.4263 to 0.8578, Indels: from 0.8069 or 0.6686 to 0.8444). Therefore, the Clair3-MP model was adjusted to include a new feature about whether a variant candidate is in a difficult region. The adjusted Clair3-MP model resulted in a small improvement (SNPs: from 0.9984 to 0.9985, Indels: 0.9736 to 0.9745), showing that the indication of difficult regions further enhances variant-calling predictions. Our experiments containing data from PacBio revealed that variant-calling performance is high in most cases, with F1 scores of at least 0.9957 when using Clair3-MP with PacBio as part of the multi-platform data or Clair3 with PacBio data only. We also found that there was an improvement in Indels calling with additional Illumina data in Clair3-MP with PacBio data (Clair3 with PacBio: 0.9933, Clair3-MP with PacBio and Illumina: 0.9969). Our work found improvements in using multi-platform data in deep-learning neural networks for variant calling. Researchers are encouraged to use Clair3-MP when using sequencing data from different platforms for variant calling and to be aware of the efficiency of using multi-platforms under various coverage and genomic contexts.

## Results

To find the best strategies for variant calling with multi-platform data, we performed whole-genome variant-calling experiments with various coverage of different combinations of Illumina, Pacbio, and ONT sequencing data downloaded from Genome-In-A-Bottle (GIAB) (10). We compared the results between Clair3 and Clair3-MP, and Clair3 call variants from single-platform data and Clair3-MP call variants from multi-platform data. The evaluation was also stratified to different genomic regions for deeper insights into variant-calling performance. We also benchmarked the impact of adding the stratification information into a neural network in this section. All running commands are available in the Supplementary Note.

### Data description

We conducted the experiments with the datasets collected in the GIAB Ashkenazi Jewish trio (HG002-child, HG003-father, and HG004-mother). The ONT sequencing data was obtained from the Human Pangenome Reference Consortium (HPRC) (11) with r9.4.1 chemistry and PromthION flowcell, with high coverage in three samples, HG002 (∼65x), HG003 (∼77x), and HG004 (∼80x), which were base-called via Guppy5. The Illumina sequencing data, obtained from precisionFDA (3), were sequenced on the NovaSeq 6000 System with a 2×151 bp high coverage PCR-free library. All three samples had coverage of 35x. The PacBio sequencing data was also from precisionFDA (3) from the Sequel II System with 2.0 chemistry and coverage of 35x for all three samples. The BAM files were obtained using the respective alignment software based on the platform of the sequencing data. Minimap2 (12) was applied to ONT data. Burrows-Wheeler Aligner (BWA)(13) was used for Illumina data, and pbmm2 (12) was used for PacBio data. All alignments used the GRCh38 human reference build. The commands for BAM files generation can be found in the Supplementary Note. For the Clair3-MP models, each multi-platform combination has its own trained model. For training each multi-platform model, we used the nine following coverages: 10x-10x, 10x-20x, 10x-30x, 20x-10x, 20x-20x, 20x-30x, 30x-10x, 30x-20x, and 30x-30x. We designed the training data in various coverage combinations to establish a model for improving robustness. HG002 and HG004 samples were used for training the Clair3-MP model. HG003 data were preserved for testing and were downsampled to 10x, 20x, and 30x. Downsampling operations for both training and testing datasets were carried out using Samtools (14). The related commands can also be found in the Supplementary Note. We downloaded the GIAB benchmark variant set version 4.2.1 as the ground truth variant sets for HG002, HG003, and HG004. The training and benchmark were constrained in the high-confidence regions also provided by the GIAB 4.2.1 benchmark set for each GIAB sample.

### Evaluation method

The results obtained by Clair3-MP on multi-platform data and Clair3 on single-platform data were evaluated using the Illumina’s Haplotype Comparison Tools (aka. hap.py) (15). The code and parameters of hap.py can be found at https://github.com/Illumina/hap.py. We used Precision, Recall, and F1-score metrics to assess the variant-calling performance. The results included performances at SNP and Indel and were stratified based on the genome stratification (version v3.1) defined by GIAB (15).

### Performance on ONT-Illumina

First, we benchmarked the Clair3-MP performance using both ONT and Illumina data with single platform data via Clair3. Fig. 2 shows the overall performance of variant calling using Clair3-MP with ONT-Illumina data, Clair3 with Illumina data or ONT data at different coverages. Fig. 2C indicates that with 30x coverage of ONT as part of the multi-platform data, the variant-calling performance was better than using either ONT or Illumina data at any coverage. The F1 scores of variant calling using Clair3-MP with multi-platform data that contains 30x ONT and Illumina data at 10x, 20x, and 30x, were 0.9880, 0.9938, and 0.9947, respectively (Table 1). However, the F1 scores of variant calling using Clair3 with Illumina data at 10x, 20x, and 30x were 0.9335, 0.988, and 0.9937, while the F1 scores using Clair3 with ONT data at 10x, 20x, and 30x were 0.9301, 0.9631 and 0.9696, respectively. The improvement in variant calling demonstrates the capability of Clair3-MP to combine ONT and Illumina data for higher performance. On the other hand, a decrease in the performance for variant calling that contains less ONT data suggests that lower-coverage ONT data might introduce background noise to calling, as there could be less or even no read support for variant candidates (Figs. 2A and 2B).

**Table 1.** The overall performance of variant calling using single- and multi-platform data (ONT and Illumina).

|  | Caller | ONT Coverage | Illumina Coverage | Precision | Recall | F1 score |
| --- | --- | --- | --- | --- | --- | --- |
| Single-platform data (ONT or Illumina) | Clair3 | 10x | / | 0.9605 | 0.9016 | 0.9301 |
|  |  | 20x | / | 0.9816 | 0.9453 | 0.9631 |
|  |  | 30x | / | 0.9857 | 0.9541 | 0.9696 |
|  |  | / | 10x | 0.9240 | 0.9432 | 0.9335 |
|  |  | / | 20x | 0.9893 | 0.9867 | 0.9880 |
|  |  | / | 30x | 0.9950 | 0.9923 | 0.9937 |
| Multi-platform data (ONT-Illumina) | Clair3-MP | 10x | 10x | 0.9822 | 0.9609 | 0.9714 |
|  |  | 10x | 20x | 0.9852 | 0.9725 | 0.9788 |
|  |  | 10x | 30x | 0.9860 | 0.9739 | 0.9799 |
|  |  | 20x | 10x | 0.9910 | 0.9800 | 0.9855 |
|  |  | 20x | 20x | 0.9938 | 0.9894 | 0.9916 |
|  |  | 20x | 30x | 0.9946 | 0.9906 | 0.9926 |
|  |  | 30x | 10x | 0.9931 | 0.9830 | 0.9880 |
|  |  | 30x | 20x | 0.9959 | 0.9917 | 0.9938 |
|  |  | 30x | 30x | 0.9966 | 0.9929 | 0.9947 |

The table shows the precision, recall and F1 score of variant calling using single-platform and multi-platform data at different coverage.

**Fig. 2.**
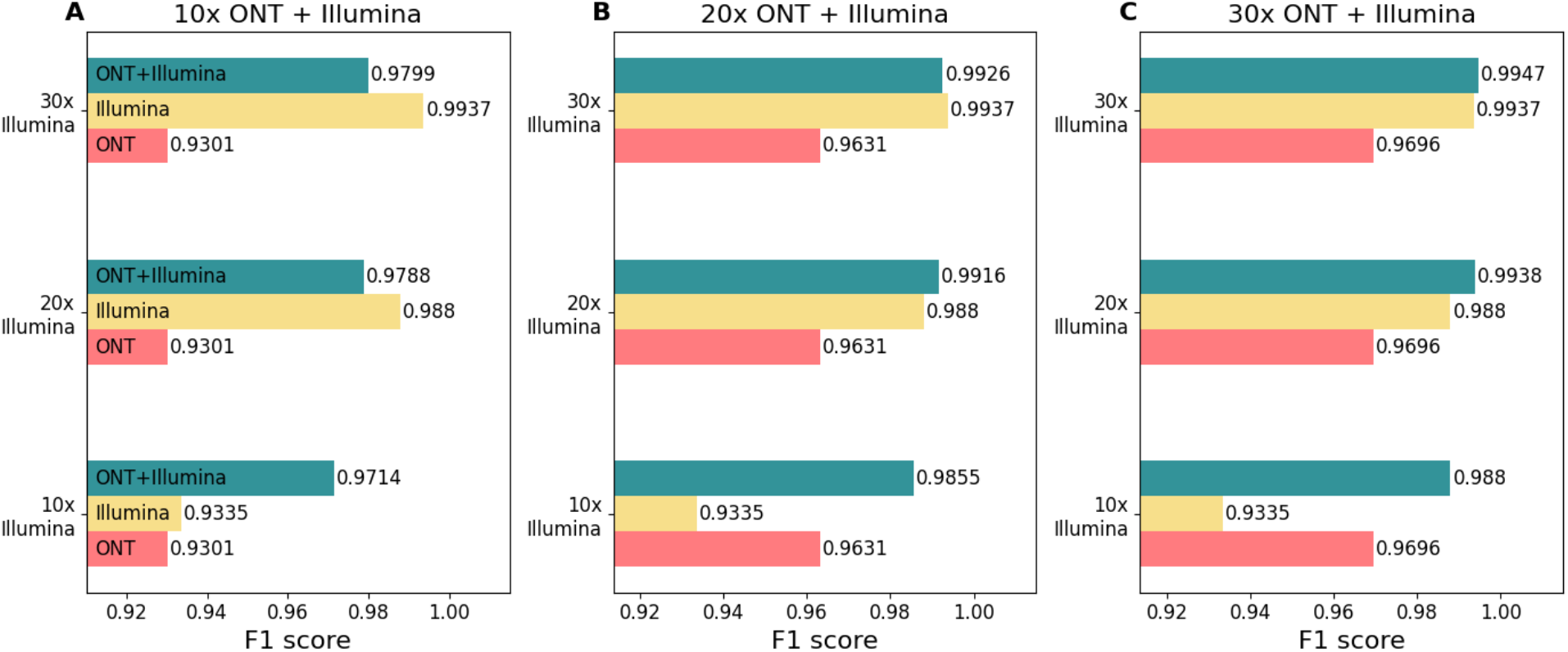
Overall F1 scores for variant calling using ONT-Illumina and single-platform sequencing datasets at different coverage. A) 10x ONT and Illumina at 10x, 20x or 30x. B) 20x ONT and Illumina at 10x, 20x or 30x. C) 30x ONT and Illumina at 10x, 20x or 30x. Each subplot contains different coverage of ONT in the multi-platform data. The x-axis is the F1 score. The y-axis is the coverage of Illumina contained in the multi-platform data. The yellow row represents the F1 score of variant calling using only Illumina data with the coverage labeled on the y-axis. The red row represents the F1 score of variant calling using only ONT data with the coverage indicated by the subplot name. The green row indicates the F1 score of variant calling using ONT and Illumina data, together with the coverage indicated by the y-axis labels and subplot names.

To investigate the impact of ONT or Illumina data as additional data for variant calling, we further present the benchmark results as a marginal graph and divide the results at the SNP and Indel level. Fig. 3 shows marginal graphs of the F1 score of variant calling comparing Clair3 with single-platform data and Clair3-MP with ONT-Illumina data at different coverage by variant type. As Fig. 3A, we concluded that with insufficient coverage of Illumina data (i.e., less than 15x), additional 10x or more ONT data significantly improves variant-calling performance. With sufficient coverage of Illumina data (i.e., more than 15x), the additional ONT data is no longer favorable for variant calling unless there is relatively high coverage of ONT (i.e., 20x). An additional 10x ONT even diminishes the performance due to the quality of the variant candidates generated by 10x ONT data. However, SNPs and Indel have different performance trends (Figs. 3B and 3C). The addition of more than 10x ONT data to any coverage of Illumina results in improved SNP-calling performance. On the other hand, we observed an improvement in Indel calling when there was insufficient coverage of Illumina data and high coverage of ONT data. When there is more than 15x Illumina data, additional ONT data at any coverage decreases the Indel-calling performance. The different trends between SNPs and Indel calling suggest that the additional cover for the genome by long ONT reads can increase SNP-calling performance (Fig. 4), while the Indel errors in ONT data, even with high coverage, affect Indel calling in multi-platform variant calling. Fig. 4 shows the alignments in the Integrative Genomics Viewer (IGV) (16) that contain true SNP cases, showing that ONT reads provide a broader range of cover for the regions that Illumina reads have trouble being mapped to.

**Fig. 3.**
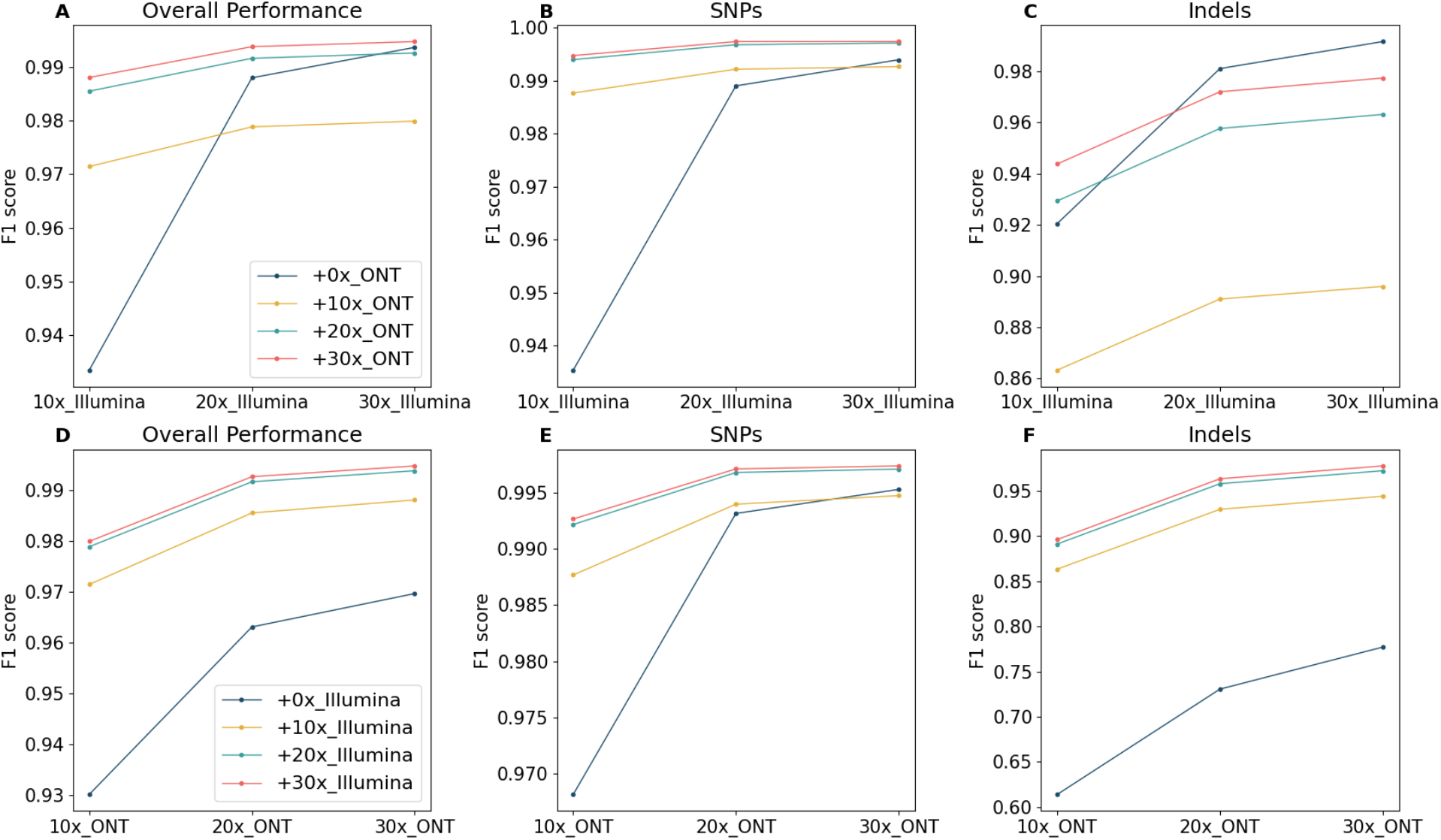
Marginal graph of the F1 score using single-platform data and ONT-Illumina data at different coverage. A) Overall F1 score in a comparison between Illumina data and ONT-Illumina data. B) SNP-calling F1 score in a comparison between Illumina data and ONT-Illumina data. C) Indel-calling F1 score in a comparison between Illumina data and ONT-Illumina data. D) Overall F1 score in a comparison between ONT data and ONT-Illumina data. E) SNP-calling F1 score in a comparison between ONT data and ONT-Illumina data. F) Indel-calling F1 score in a comparison between ONT data and ONT-Illumina data. The x-axis indicates the coverage of a single-platform data included in the variant calling for combining and not combining the various coverage of data from the other platform. Each trend line indicates the coverage of the other platform data added in the variant calling in Clair3-MP. The intersections of the trend lines reveal the optimal case for combining ONT and Illumina data for variant calling.

**Fig. 4.**
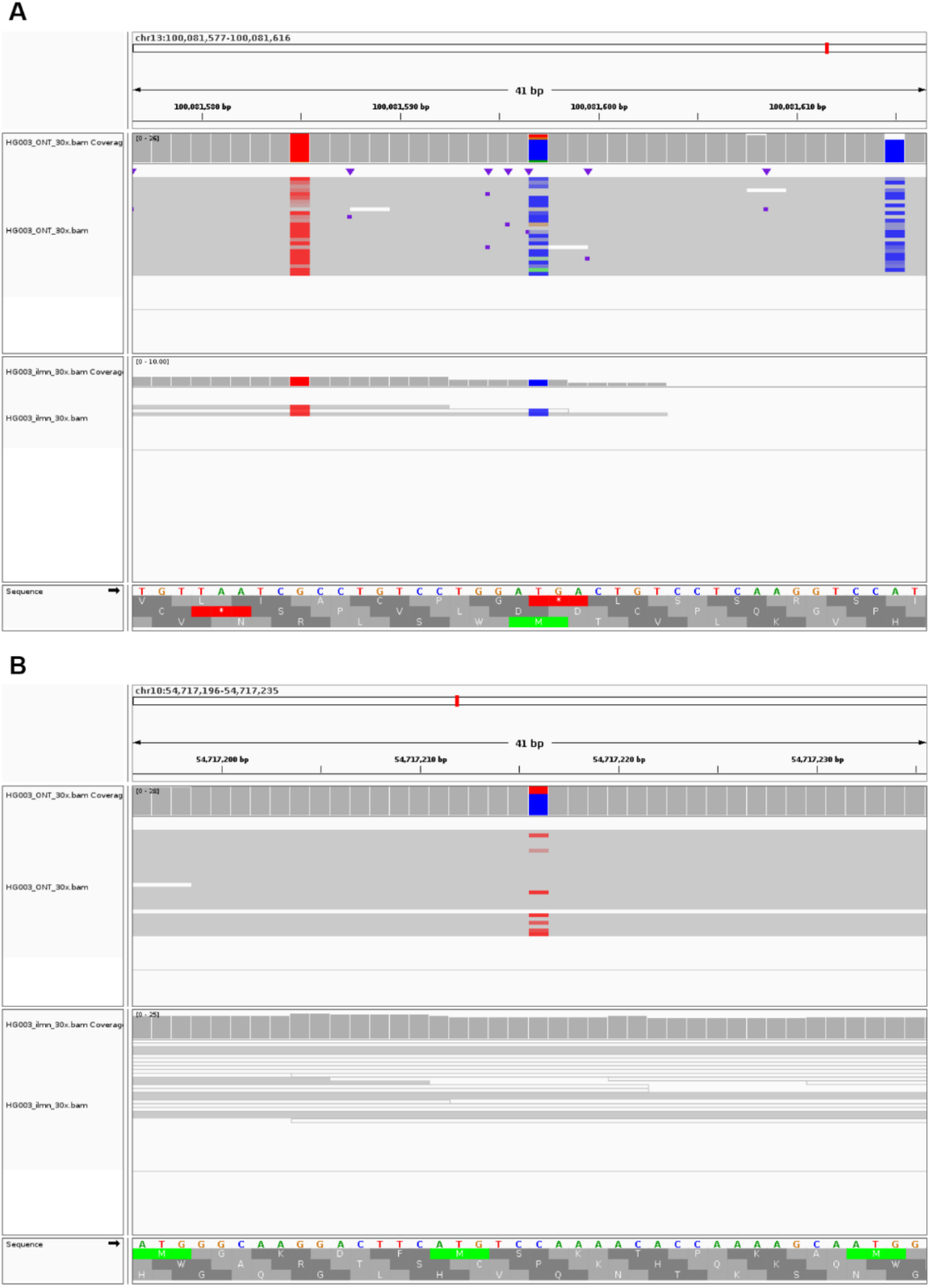
Two true variants called by Clair3-MP with ONT and Illumina (both at 30x) A) A true SNP that was called using ONT-Illumina data in Clair3-MP but not called using Illumina data in Clair3. This variant is at chr13:100081597; it is a homozygous T-to-C substitution. The region is not mapped by Illumina reads, even though there are reads supporting the allele sequence. By covering the regions that Illumina reads have trouble being aligned to, with ONT’s long reads, ONT-Illumina data increases the performance of SNP calling with Clair3-MP. B) A true SNP that was called using ONT-Illumina data in Clair3-MP but not called using either ONT or Illumina data in Clair3. This variant is at chr10:54717216; it is a heterozygous C-to-T substitution. The region is mapped by ambiguous Illumina reads with mapping quality of 0, meaning that these reads can be mapped to other regions too. However, with only ONT data, there is not enough support for prediction. The variant was called when combining ONT and Illumina data, showing the capability of Clair3-MP to recover true variants that are not called using only ONT or Illumina data.

Figs. 3D, 3E, and 3F show line charts of the F1 score of variant calling comparing Clair3 with ONT data and Clair3-MP with ONT-Illumina data at different coverage by variant type. Other than in SNP calling using sufficient coverage of ONT data (i.e., 30x) with low coverage of Illumina data (i.e., 10x), any additional Illumina data can increase the variant-calling performance using only ONT data. This indicates that additional and accurate Illumina reads can improve calling by providing strong and positive signals, especially in Indel calling, unless the Illumina data is at such low coverage that it decreases the SNP calling performed by high coverage of ONT data.

### Performance of various stratification regions on ONT-Illumina

We further divided the performance results for ONT-Illumina into different reference genome regions to explore the reference impact when using multi-platform data. The reference genome regions (stratification) were obtained from GIAB. There are 10 major types of stratification: Low Complexity, Functional Technically Difficult, Genome Specific, Functional Regions, GC Content, Mappability, Other Difficulties, Segmental Duplications, Union, Ancestry, and XY. The stratification analysis was conducted using a testing datasets of 30x ONT and 30x Illumina from GIAB HG003.

Fig. 5A shows SNP-calling performance with Clair3-MP combining 30x ONT and 30x Illumina data compared to using either ONT or Illumina data in Clair3 in eight stratification regions. The complete figures and tables stratifying the results of SNP calling into all the different genomic regions are in the Supplementary Figures S1 to S2 and Supplementary Tables S2 to S4. Fig. 4A shows that the SNP F1 score of Clair3-MP 30x ONT-Illumina data (large repeat regions F1 score: 0.9973, satellites regions: 0.9965, segmental duplications regions: 0.9653, segmental duplications region longer than 10K bp: 0.9605) outperformed both Clair3 with 30x ONT data (large repeat regions F1 score: 0.9963, satellites regions: 0.9936, segmental duplication regions: 0.9565, segmental duplication regions longer than 10K bp: 0.9501) and Clair3 with 30x Illumina data (large repeat regions F1 score: 0.9844, satellites regions: 0.9861, segmental duplications regions: 0.9177, segmental duplications regions longer than 10K bp: 0.9087). The SNP F1 scores using either ONT or Illumina data with Clair3 have very similar values in the regions with GC content that is less than 30% or greater than 55%, which are 0.9947 and 0.9951. This shows that this region is well-covered by either ONT or Illumina and that SNP calling reaches a relatively high performance with Clair3. When ONT and Illumina are combined with Clair3-MP, the performance can be improved to 0.9975. This suggests that the integration of ONT and Illumina data in Clair3-MP can enhance SNP calling in regions using either data that already achieved relatively high performance. Also, compared to using only Illumina data for variant calling, Clair3-MP with ONT-Illumina data provides extra coverage for the genome from ONT’s long reads and leverages the strength of both datasets to further improve performance, even in difficult regions.

**Fig. 5.**
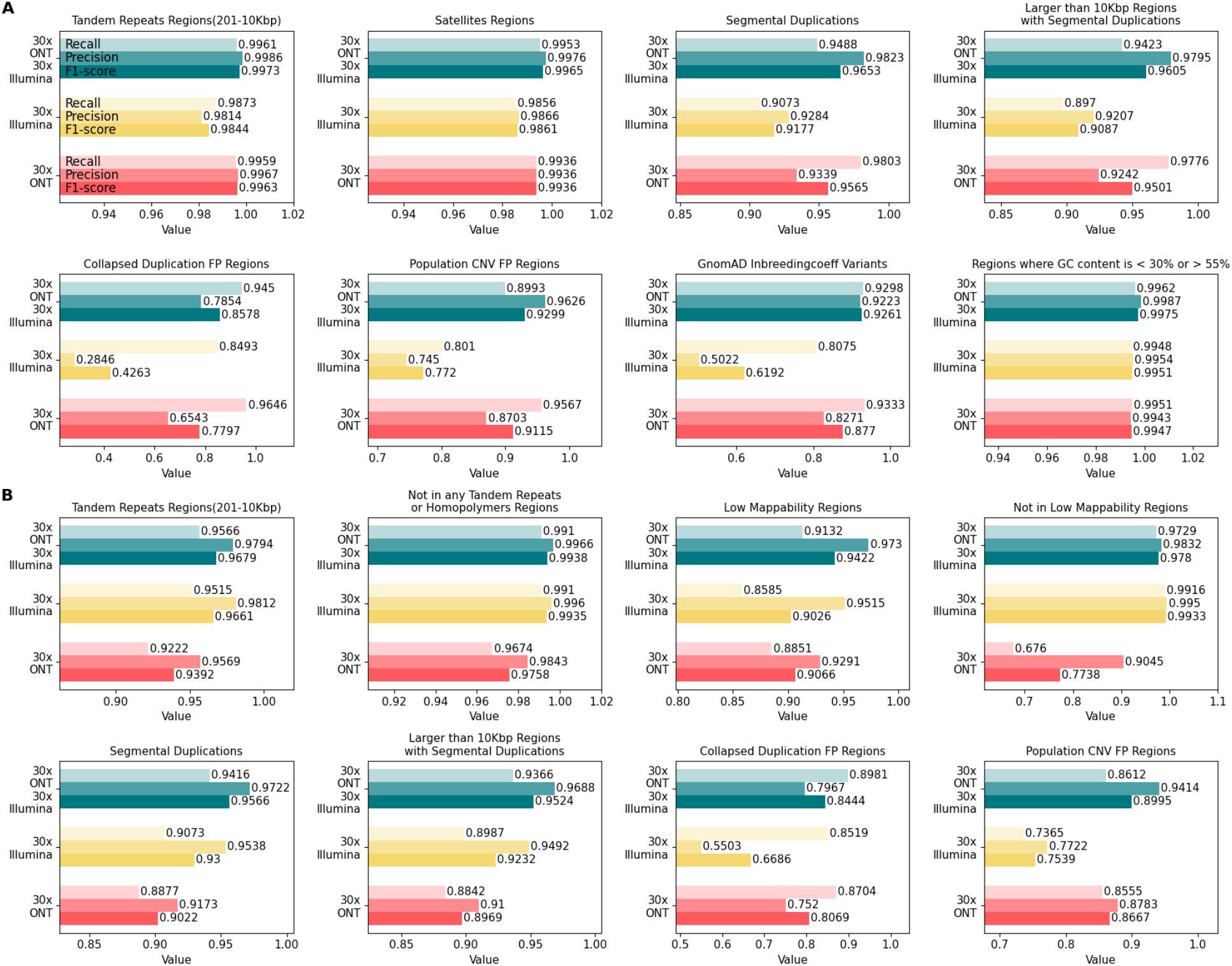
The performance in stratifications for 30x ONT-Illumina data in comparison with ONT and Illumina data. A) SNP-calling performance in eight stratifications. B) Indel-calling performance in eight stratifications. The x-axis is the value for precision, recall and F1 score. The y-axis is the label for data used in the variant calling.

On the other hand, we also found a list of regions that are challenging for Clair3-MP. Collapsed duplication FP regions and population CNV FP regions are both related to collapsed duplications errors in the GRCh38, recently identified and corrected by the T2T consortium (17). These regions are populated with paralog-specific variants (PSV) – variation among paralogous sequences, which impact short-read variant calling (18). In addition, the long segmental duplications with high sequence identity are a challenge for long-read mapping accuracy (18). Therefore, we found lower SNPs F1 scores using only ONT or Illumina data (collapsed duplication FP regions for either ONT or Illumina: 0.7797, 0.4263; population CNV FP regions: 0.9115, 0.7720). Although challenges for ONT and Illumina reads vary in this region, the combination of ONT and Illumina data demonstrates great improvement in SNP F1 scores (collapsed duplication FP regions: 0.8578; population CNV FP regions: 0.9299). Moreover, other difficult regions that are enriched by variants with excess heterozygosity, which are defined by the InbreedingCoeff from the gnomAD database, were found to contain numerous false positives in variant calling (17). This is also reflected by our benchmark – the precision for SNP calling with only Illumina data is 0.5022. Like the collapsed duplication regions, the precision of SNP calling using ONT-Illumina data was increased to 0.9223 in this region, leading to an improvement in the F1 score from 0.6192 to 0.9261. This shows the capability of multi-platform data to improve variant-calling performance in difficult regions that are not accurately called by either single-platform dataset.

In contrast, Indel calling using both data sets decreased for most of the genomic regions due to the high indel error rate in ONT reads (Supplementary Figures S3 to S4 and Supplementary Tables S2 to S4). However, similar to the improvement for SNP calling, Fig. 5B shows that the indels F1 score of Clair3-MP 30x ONT-Illumina data (large repeat regions F1-score: 0.9679, segmental duplication regions: 0.9566, segmental duplication regions longer than 10K bp: 0.9524, collapsed duplication regions: 0.8444, population CNV FP regions: 0.8995) outperformed both Clair3 with 30x ONT data (large repeat regions F1 score: 0.9392, segmental duplication regions: 0.9022, segmental duplication regions longer than 10K bp: 0.8969, collapsed duplication regions: 0.8069, population CNV FP regions: 0.8667) and Clair3 with 30x Illumina data (large repeat regions F1 score: 0.9661, segmental duplication regions: 0.9300, segmental duplication regions longer than 10K bp: 0.9232, collapsed duplication regions: 0.6686, population CNV FP regions: 0.7539). In addition, we found improvement in the indel F1 score in low mappability regions (Clair3 with ONT: 0.9066; Clair3 with Illumina: 0.9026; Clair3-MP with ONT-Illumina: 0.9422). The improvement of Indel calling in these regions indicates that ONT and Illumina data can compensate for each other and that ONT reads do not introduce misleading signals for incorrect prediction. More importantly, in the regions that do not overlap with any tandem repeats or homopolymers, the ONT-Illumina data results in a slight improvement in the indel F1 score (Clair3 with ONT: 0.9758; Clair3 with Illumina: 0.9935; Clair3-MP with ONT-Illumina: 0.9938). This hints that the noise/misleading signals of Indel calling caused by the Indel error rate lie in the regions with tandem repeats and homopolymers. Although there are regions with a higher F1 score using Clair3-MP with ONT-Illumina data in indel calling, these regions account for only small parts of the genome, and the improvement cannot compensate for the lower performance in the other genomic regions.

### Performance with the addition of stratification information to Clair3-MP

This section describes the benchmarking results of adding stratification information as a feature to the Clair3-MP full-alignment model. Based on the “Stratification on ONT-Illumina” results, we observed that the performance of Clair3-MP with ONT-Illumina data varies across different genomic regions and improved in difficult regions. We investigated whether using various genomic information for multiplatform data can further improve variant-calling performance. We integrated the stratification information, which is interpreted as whether the variant candidate is in a difficult genomic region defined by the GIAB BED file, into Clair3-MP and trained a new ONT-Illumina model from GIAB HG002 data. We compared the newly trained model (with stratification information) with the original Clair3-MP model and tested their performance at the sequencing data of 30x ONT and 30x Illumina from HG003 chromosome 20, which has always been excluded from model training. Fig. 6 is the F1 score of SNPs and Indel calling using the original Clair3-MP model compared to the Clair3-MP model with the stratification information added. It shows that including the information about whether the variant candidate is in a difficult genomic region for prediction yields comparable results. Adding the stratification increases the performance of SNPs (original Clair3-MP: 0.9984, new Clair3-MP: 0.9985) and Indel calling in Clair3-MP (original Clair3-MP: 0.9736, new Clair3-MP: 0.9745). This small but measurable improvement provides the important message that if a variant candidate is in a difficult region, Clair3-MP not only combines the ONT and Illumina data, but also allocates them based on the difficulty of the region, for best variant-calling performance.

**Fig. 6.**
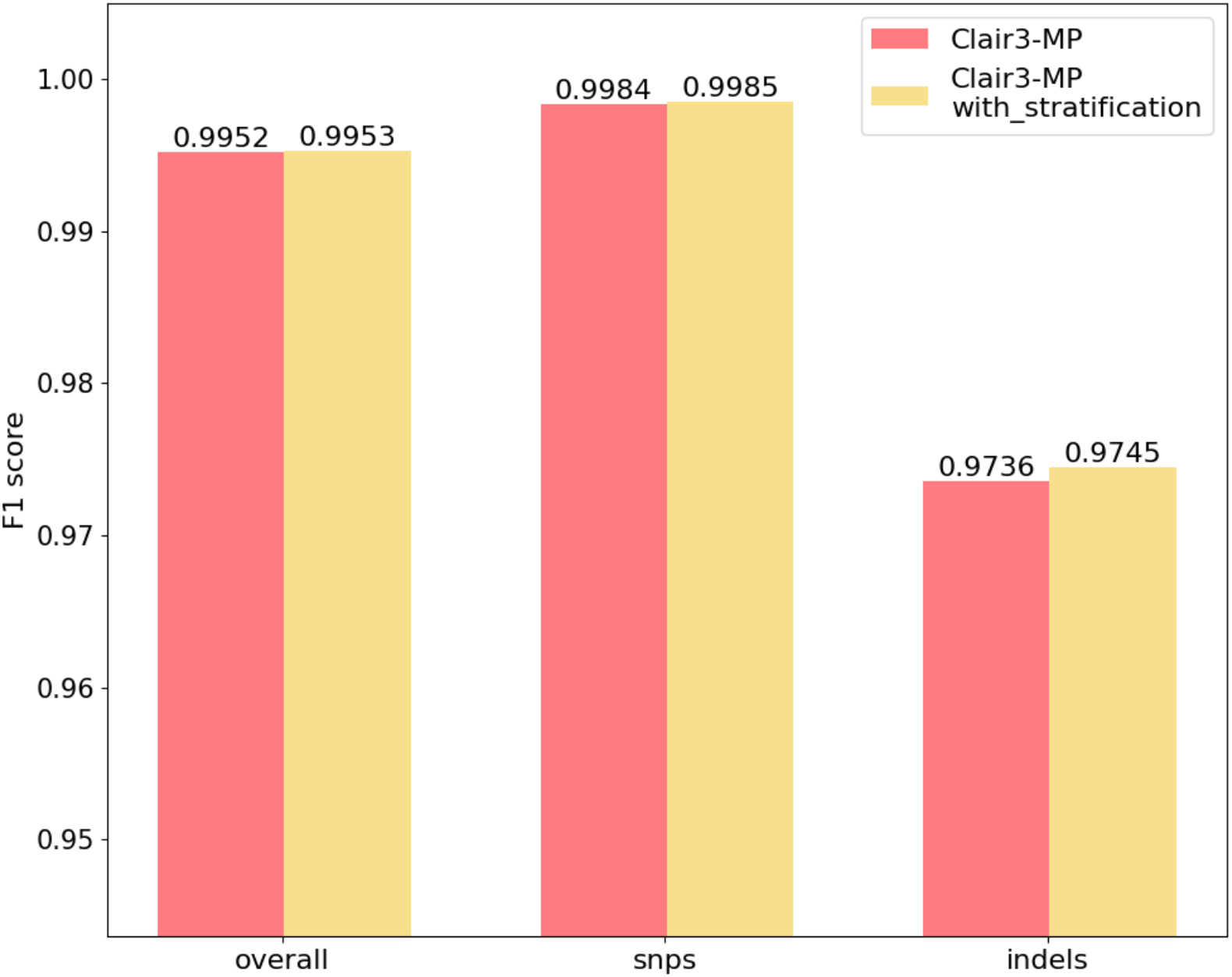
The F1 score for variant calling, original Clair3-MP model v.s.Clair3-MP model with the stratification channel. The red column is the F1 score for variant calling using the original Clair3-MP model, and the yellow column indicates the F1 score for variant calling using the Clair3-MP model trained with the stratification information.

### Performance on ONT-PacBio and PacBio-Illumina

In this section, we present variant-calling performance using Clair3-MP with multi-platform data involving PacBio data. Featuring long- and high-read accuracy, PacBio was benchmarked with a high F1 score (>0.99) in variant calling (3). We explored the effect of using PacBio data as a part of the data within a multi-platform framework. We trained two new models for ONT-PacBio and PacBio-Illumina data and tested the performance with HG003 data. We compared the performance of the newly trained models with Clair3. The execution commands are available in the Supplementary Note. Fig. 7 is a column chart that shows the F1 score in variant calling in three areas – overall performance, SNP calling and Indel calling – with different data combinations. We found that Clair3-MP with PacBio and other platform data achieved results comparable to Clair3 with PacBio data. Variant-calling performance was high, with at least a 0.9957 F1 score in most cases when PacBio was part of the multiplatform data or as single-platform data. Of note, Indel calling using Clair3-MP with PacBio-Illumina data had an F1 score of 0.9969, which was higher than 0.9933 using Clair3 with PacBio data. This suggests that due to the higher indel error rate in PacBio data compared to Illumina data, additional sufficient and accurate Illumina data containing fewer indel errors can improve the performance in Indel calling with Clair3-MP, but not in SNP calling. However, Clair3-MP with ONT-PacBio had a worse performance in variant calling than Clair3 with PacBio (PacBio: 0.9990, ONT-PacBio: 0.9986) and Indel calling (PacBio: 0.9933, ONT-PacBio: 0.9771). The decrease in indel-calling performance echoes back to the impact of ONT discussed earlier, whereby the indel error rate affected Indel calling in multi-platform data. The slight decrease in SNP calling suggests that Clair3-MP still has room for improvement to eliminate noise from the ONT reads.

**Fig. 7.**
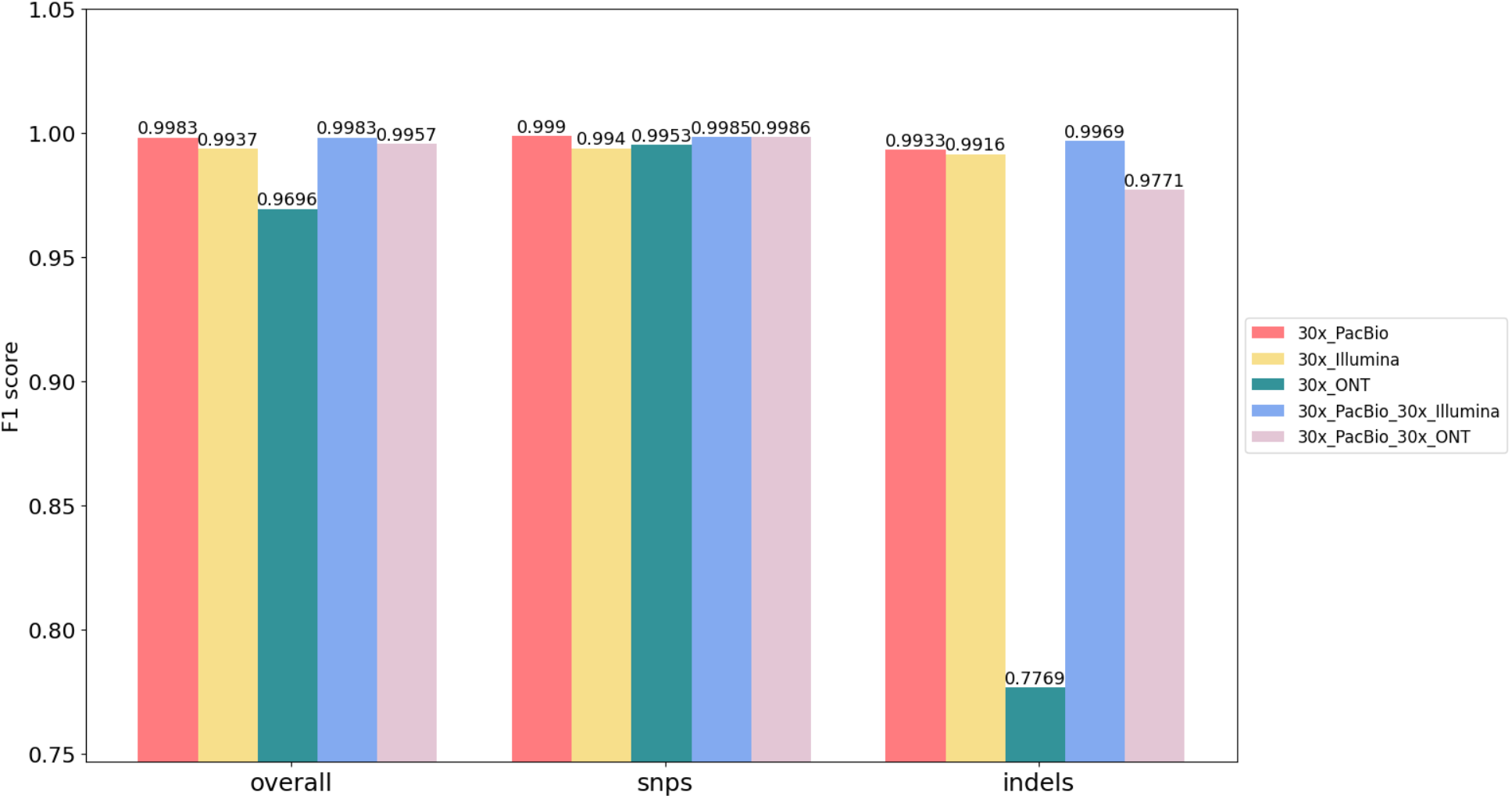
The F1 score of variant calling using multi-platform data involving the PacBio and single-platform datasets. The red, yellow, and green columns indicate the F1 score of variant calling using only 30x PacBio data, 30x Illumina data, and 30x ONT data, respectively. The blue column shows the F1 score of variant calling combining 30x PacBio and 30x Illumina data in Clair3-MP, and the pink column is the F1 score using 30x PacBio and 30x ONT data.

## Discussion

Through a series of extensive experimental investigations conducted in diverse scenarios, we demonstrated that the utilization of multi-platform data with Clair3-MP leads to a notable improvement in variant calling. Our results show that Clair3-MP exhibits robust performance across datasets of varying error profiles and coverage. By stratifying the variant calling results from Clair3-MP with ONT-Illumina data (both at 30x), we also found that an even higher performance in SNP calling by Clair3-MP with ONT-Illumina arises primarily from an improvement in large tandem repeat regions, satellite regions, segmental duplication regions, collapsed duplication regions, and GC content regions. The improvement shows that Clair3-MP can leverage the strength of both data sets to reach higher performance. Although overall Indel calling using Clair3-MP with ONT-Illumina data is affected by noise introduced by a higher indel error rate in the ONT reads, we found that Clair3-MP achieves better Indel calling in the large tandem repeat regions, low Illumina-read mappability regions, and segmental duplication and collapsed duplication regions. Based on our stratification analysis, we concluded that Clari3-MP with ONT-Illumina data can attain better variant-calling performance in challenging genomic regions than Clair3 with ONT or Illumina data can. Given this conclusion, we adjusted the Clair3-MP model to include information about whether a variant candidate is in a difficult region to further enhance the variant-calling performance. Adding stratification improves the capability of Clair3-MP with ONT-Illumina data for variant calling in the new model, and the capability to leverage both datasets can be adjusted based on the genomic region. The high performance in variant calling by Clair3 with PacBio data prompted us to investigate whether the inclusion of data from additional platforms could yield even better performance in Clair3-MP. We ensured that the combinations used in Clair3-MP provided sufficient data coverage. We observed that adding 30x Illumina data to 30x PacBio data in Clair3-MP resulted in higher performance in Indel calling than Clair3 with PacBio data. However, the impact of Illumina and ONT reads on SNP calling, and overall variant calling in ONT-PacBio data highlights the need for further refinement of the Clair3-MP model to mitigate noise from these sources.

The improvement in variant calling using Clair3-MP with ONT-Illumina validates the hypothesis that multi-platform data can leverage the strengths of different sequencing data. It has been previously discussed that for high coverage of ONT data, SNP calling demonstrated improved performance with developed methods due to its broader range of cover for the genome. Therefore, the additional 30x ONT data to Illumina data in Clair3-MP can compensate for the limitations of Illumina data for SNP calling, resulting in an improvement in overall calling performance. However, with additional ONT data of lower than 20x, the performance for variant calling using Clair3-MP with ONT-Illumina is not as improved as with the additional 30x ONT data. This indicates the relevance of coverage for the ONT data to variant-calling improvement in Clair3-MP. ONT data with lower coverage lacks read support for the allele sequence (Supplementary Figure S5A) and even for the region (Supplementary Figure S5B), as it introduces background noise in the neural network. Moreover, the decreased indel-calling performance in most of the coverage combinations, even with 30x ONT data, shows that the high indel error rate of ONT reads adversely impacts variant calling in Clair3-MP with ONT-Illumina data. We provide examples in Supplementary Figure S6 to illustrate how the presence of multiple insertion sequences supported by ONT data can introduce ambiguity to Illumina read signals, thereby affecting the quality of variant candidates. The histogram in Supplementary Figure S7 indicates how the quality scores have shifted in the indels called by all Clair3 with Illumina, Clair3 with ONT, and Clair3-MP with both data. The quality scores of indels from Clair3-MP show a decreasing trend compared to Clair3 with Illumina data, providing an overall understanding of how ambiguity among ONT data impacts the quality of indel calling. With the above conclusions and the message conveyed by Fig. 3, we recommend that researchers perform variant calling using Clair3-MP with ONT-Illumina data with at least 30x coverage of ONT data. If there is low coverage of Illumina data, additional ONT data at any coverage also guarantees improvement in variant calling with Clair3-MP. Moreover, the acquisition of additional Illumina data at any coverage level can be useful when only ONT data is available, as it has been confirmed to improve variant-calling performance.

The stratification analysis provides valuable insights into variant calling using ONT-Illumina data. It is reasonable to believe that multi-platform data can leverage the strengths of the data and complement the weaknesses of each dataset. Through our stratification analysis, we found that the improvement in variant calling with Clair3-MP lies primarily in the difficult regions, which include the regions in which Illumina reads are susceptible to mapping and duplication resulting from incorrect mapping due to PSVs. Fig. 8 shows two cases of true variants in the difficult regions of low mappability and collapse duplication. Fig. 8A is a true SNP in the collapse duplication regions mapped by misaligned Illumina reads due to the PSA in the duplication sequences, and Fig. 8B is a true variant in a low mappability region, to which only ambiguous Illumina reads can map. In these cases, Clair3-MP leverages the strengths of confident variant candidates from the ONT data to compensate for the ambiguity in Illumina reads. In the regions in which Illumina data have sufficient coverage and high mapping quality, and in which variant calling in Clair3 with only Illumina reads already provides adequate performance, the additional ONT data leads to a decrease in Indel calling performance, such as smaller low complexity regions shorter than 200 bp (Supplementary Figure S4). Stratifying the performance of multi-platform data for variant calling in different genomic contexts allows researchers to make informed decisions regarding the use of multiplatform data to achieve optimal performance in their area of interest.

**Fig. 8.**
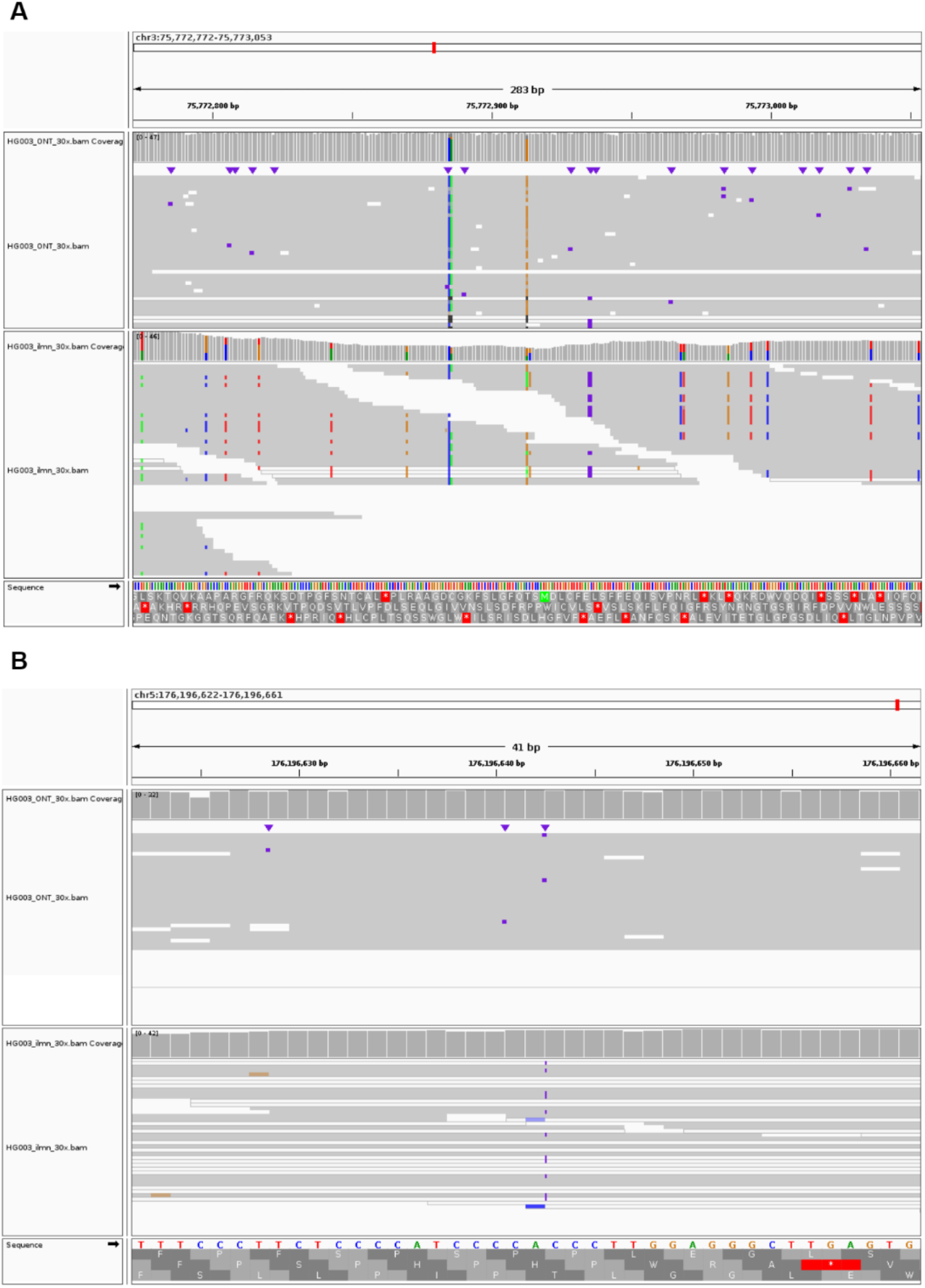
Two true variants in difficult regions, that were called using ONT-Illumina data in Clair3-MP. A) This variant is at chr3:75772913; it is a homozygous C-to-G substitution in the collapse duplication region. The Illumina reads are mapped incorrectly in this region due to PSVs, therefore this variant were not called only using Illumina data, but the ONT read can cover the region. Variant calling in this region with Illumina reads is problematic, so the addition of ONT data helps recover the SNP. B) This variant is at chr5: 176196642; it is a heterozygous C insertion in a low-mappability region. The region is mapped by ambiguous Illumina reads with mapping quality of 0, meaning that these reads, even with insertion sequences, can be mapped to other regions confidently as well.

The stratifying results clearly demonstrate the capability of Clair3-MP to improve variant calling in difficult genomic regions. To further leverage the strengths of multi-platform data, we adjusted the Clair3-MP model to incorporate information on whether a variant candidate is in a challenging genomic region. The success in improving variant-calling performance hints that a deep-learning network can take into account the genomic context information for higher accuracy. Fig. 9 shows the feature activation of a true variant called using only the adjusted Clair3-MP model. No features in the ONT or Illumina reads were activated in the Clair3-MP model, but the features in the Illumina reads were activated intensely in the Clair3-MP stratification model. The re-activation of features suggests that the inclusion of stratification information allows the network to recover true SNPs that were previously missed in the multi-platform data.

**Fig. 9.**
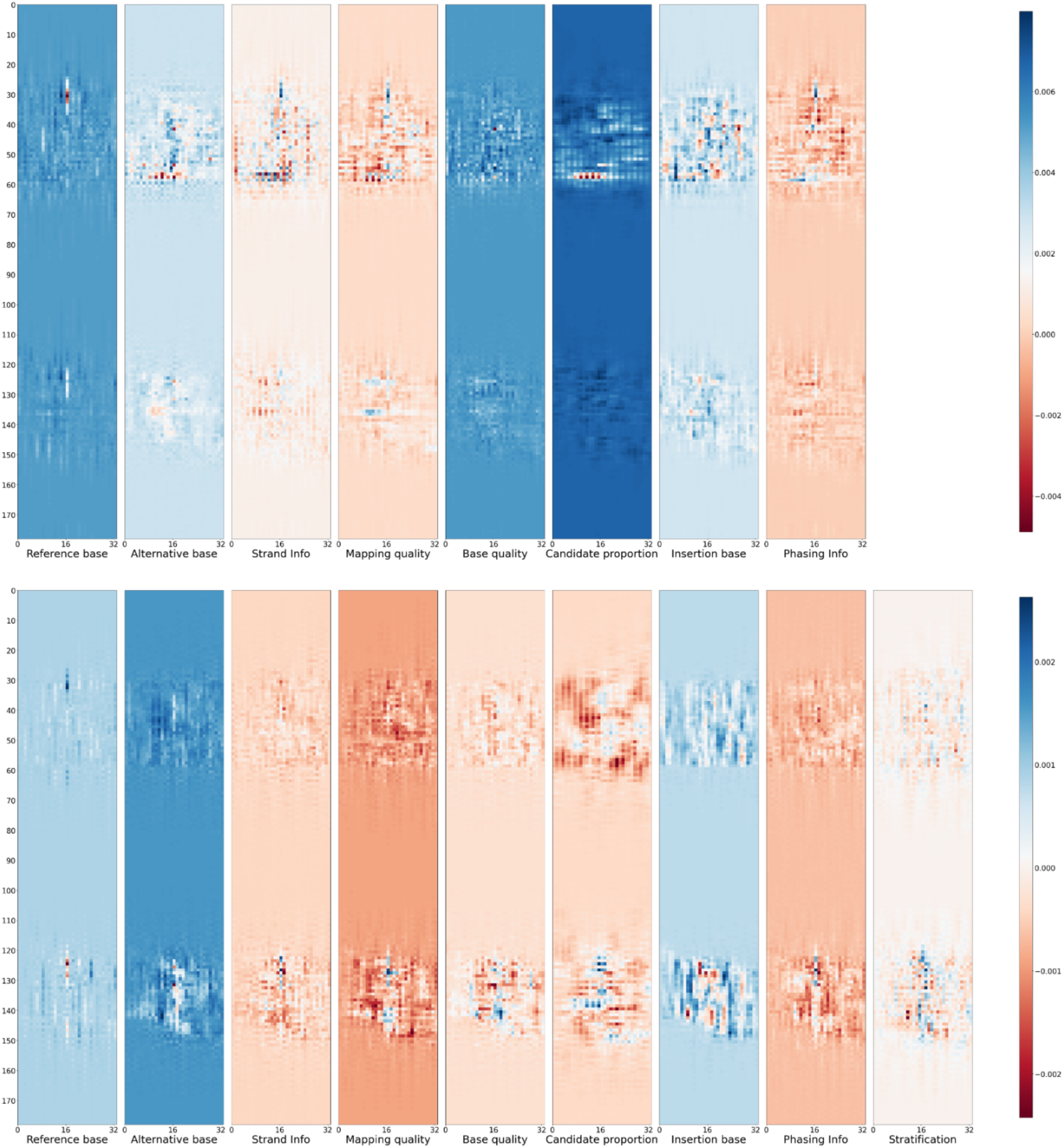
Activation graph of a true variant. This variant was called by the Clair3-MP model with the stratification feature but was not called in the original Clair3-MP model. The top rows of both graphs are the ONT reads, and the bottom rows are the Illumina reads.

In our study, we trained the PacBio-involved model to illustrate multi-platform effects when including the PacBio data in Clair3-MP. Our findings demonstrate that the inclusion of data from other platforms with PacBio data leads to comparable results, as Clair3 with PacBio achieved high performance. Therefore, we did not train a three-platform model.

There are some challenges, and future work remains to be done to utilize multiple platform data. We benchmarked Clair3-MP only on whole genome sequencing (WGS) data. Other experiments combined both target region sequencing (like sequencing high-sequencing coverage at targeted sequencing regions) and WGS (like using ONT or PacBio WGS). These designs challenge Clair3-MP for its unbalanced data coverage across the reference genome. In our future research, we aim to enhance the versatility of Clair3-MP by enabling the automatic acceptance of unbalanced data, such as Illumina-only and ONT-only data, in training an ONT-Illumina MP model. However, the current design of Clair3-MP showed marginal improvement in in indel calling. This is due to the high indel error rate in ONT sequencing reads, which confuses the model from converging well. Some techniques, like sequence error correction, realignment, and alignment to consensus, may further improve the calling accuracy.

## Conclusions

Our studies demonstrate the capability of combining multi-platform data for improving variant calling, and validate the strengths and weaknesses of variant calling using ONT and Illumina data, through various experiments with Clair3-MP. Clair3-MP with ONT-Illumina data outperformed Clair3 with ONT or Illumina data in difficult regions. In addition, the experiments at different coverage provide insights for research involving variant calling using sequencing data from different platforms. Combined with stratification analysis, researchers can decide what the best scenarios are for performing variant calling with multi-platform data based on the nature of the study. Building on the stratification results, we further modified the deep-learning model to better process multi-platform data based on the genomic region, resulting in improvements for both SNPs and indel calling.

## Methods

### Clair3-MP workflow

Clair3-MP has two parts for variant callings: the first part generates high-quality candidates using Clair3’s pileup models to phase the input alignments independently based on the platform; and the second part takes all the phased alignments and pileup candidates extracted prior to the first part and loads them into the Clair3-MP full-alignment model for variant calling. As Fig. 1 shows, Clair3-MP took the alignments from two different sequencing platforms of the same sample. This approach extracts pileup candidates using Samtools mpileup (14) and processes them separately in Clair3’s pileup models, which are tailored to the specific sequencing platforms. Then, the pileup calls are organized into variant calls (genotypes 0/1, 1/1, and 1/2) and reference calls (0/0), and these two groups are ranked by variant quality (QUAL). The top 70% of heterozygous (genotype 0/1) SNP calls are used to phase the variants and haplotag the alignments using WhatsHap (19). The phased alignments are then fed as input for haplotype-aware full-alignment calling. Lastly, all the pileup candidates are processed again in the designated full-alignment model trained for the platform combination to output one final variant set for this sample. To support the processing of alignments from different platform combinations, Clair3-MP trained a series of full-alignment models, including ONT-Illumina, ONT-PacBio and PacBio-Illumina platform combinations.

### Clair3-MP’s Input/output

As the pileup input is processed separately for each sequencing platform through Clair3’s pileup models, during the pileup calling, Clair3-MP utilizes the pileup input design as Clair3, which comprises 594 integers – 33 genome positions with 18 features at each position. For full-alignment input, to incorporate details from different platforms, Clair3-MP designed a comprehensive full-alignment input comprising 46,992 integers – eight channels of 33 genome positions and a maximum 178 reads on two platforms. We also attempted to integrate stratification information into full-alignment input to determine the contribution of genomic context to variant-calling performance. Supported by the full-alignment model, which adds the stratification information channel, the input comprises 52,866 integers, with nine channels of features. The output of the pileup model contains only two tasks for efficiency, 21 genotypes, and zygosity, which are also included in the output of the full-alignment models with two more indel length tasks. The details of the pileup and full-alignment features and output tasks are described in the Supplementary Note.

### Clair3-MP full-alignment model

The Clair3-MP model is an N-to-1 model (N = the number of platforms) that can input all the alignments from different sequencing platforms for the same sample and output one set of variants (20). For each platform, the full-alignment information is converted into eight different feature channels: reference base, alternative base, strand information, mapping quality, base quality, candidate proportion, insertion base, and phasing information channels. An additional stratification information channel is appended using the Clair3-MP stratification model. The information from each platform is aggregated for each channel and used as input for the Clair3-MP model.

### Training the Clair3-MP model

To train the Clair3-MP model with different platform combinations at various coverages, we first used the unified true variants from the Representation Unification module from Clair3 to phase the alignments of ONT, Illumina, and PacBio data. The phased alignments for each platform and each training sample were downsampled to 10x, 20x, and 30x for multiple coverage combinations. Then, the Clair3 pileup calling was performed for each downsampled and phased alignment. Each pileup calling candidates group was selected based on the ratio of 1:5 for variant calls and reference calls to avoid introducing a massive number of negative samples in model training. To train the model for each platform combination (i.e., ONT-Illumina, ONT-PacBio, PacBio-Illumina), we used the nine following coverages: 10x-10x, 10x-20x, 10x-30x, 20x-10x, 20x-20x, 20x-30x, 30x-10x, 30x-20x, and 30x-30x.

### Training the Clair3-MP model with stratification information

After conducting stratification analyses using the Clair3-MP pre-trained model, we found that the improvement in variant calling using Clair3-MP with ONT and Illumina data came mainly from improvements in difficult regions, which it is difficult to map accurately by the Illumina reads or contain true variants overlapped with the PSV. To account for this, an additional stratification channel was added to the Clair3-MP model, with a 50/100 value, indicating whether the variant candidate is in one of the difficult regions. The difficult regions are defined by ‘alldifficultregions.bed.gz’ from GIAB stratification v2 for the GRCh38 reference genome. This stratification contains a union of difficult genomic regions, such as all tandem repeat regions, difficult-to-map regions, and segmental duplication regions. This stratification was selected after a series of attempts to include different unions of difficulties in the model. We retrained the new Clair3-MP stratification channel from scratch. The results are shown in the section “Performance when stratification information is added to Clair3-MP”.

## List of abbreviations

Indels: Insertions and deletions
SNP: Single nucleotide polymorphism
Clair3-MP: Clair3 Multi-Platform
NGS: Next-generation sequencing
TGS: Third-generation sequencing
GATK: Genome analysis toolkit
ONT: Oxford Nanopore Technologies
PacBio: Pacific Biosciences
GIAB: Genome-in-a-bottle
BAM: Binary alignment map
HPRC: Human Pangenome Reference Consortium
BWA: Burrows-wheeler aligner
WGS: Whole genome sequencing

## Declarations

### Ethics approval and consent to participate

Not applicable.

### Consent for publication

Not applicable.

### Availability of data and materials

Project Name: Clair3-MP

Project homepage: https://github.com/HKU-BAL/Clair3-MP

Operating system: Linux

Programming languages: Python, Shell, C++, C

License: BSD 3-Clause

Data availability: The links to the reference genome, truth variant sets, benchmarking materials, and sequencing data from ONT, Illumina, and PacBio can be found in the Supplementary Note. The commands used in the study are available in the Supplementary Note. All results, including VCFs and logs, are available at http://www.bio8.cs.hku.hk/clair3_mp.

### Authors’ contributions

R. L., T. L., and J. S. conceived the study. H. Y., J. S., and Z. Z. implemented Clair3-MP. H. Y. and J. S. designed the experiments. H. Y. analyzed the results and wrote the paper. All authors evaluated the results and revised the manuscript.

## Acknowledgments

Not applicable.

## Reference

1. Olson ND, Wagner J, Dwarshuis N, Miga KH, Sedlazeck FJ, Salit M, et al. Variant calling and benchmarking in an era of complete human genome sequences. Nat Rev Genet. 2023.

2. Hassan S, Bahar R, Johan MF, Mohamed Hashim EK, Abdullah WZ, Esa E, et al. Next-Generation Sequencing (NGS) and Third-Generation Sequencing (TGS) for the Diagnosis of Thalassemia. Diagnostics (Basel). 2023;13(3).

3. Olson ND, Wagner J, McDaniel J, Stephens SH, Westreich ST, Prasanna AG, et al. PrecisionFDA Truth Challenge V2: Calling variants from short and long reads in difficult-to-map regions. Cell Genom. 2022;2(5).

4. DePristo MA, Banks E, Poplin R, Garimella KV, Maguire JR, Hartl C, et al. A framework for variation discovery and genotyping using next-generation DNA sequencing data. Nat Genet. 2011;43(5):491–8.

5. Rang FJ, Kloosterman WP, de Ridder J. From squiggle to basepair: computational approaches for improving nanopore sequencing read accuracy. Genome Biol. 2018;19(1):90.

6. Poplin R, Chang PC, Alexander D, Schwartz S, Colthurst T, Ku A, et al. A universal SNP and small-indel variant caller using deep neural networks. Nat Biotechnol. 2018;36(10):983–7.

7. Zheng Z, Li S, Su J, Leung AW-S, Lam T-W, Luo R. Symphonizing pileup and full-alignment for deep learning-based long-read variant calling. Nature Computational Science. 2022;2(12):797–803.

8. Ramachandran A, Lumetta SS, Klee EW, Chen D. HELLO: improved neural network architectures and methodologies for small variant calling. BMC Bioinformatics. 2021;22(1):404.

9. Holley G, Beyter D, Ingimundardottir H, Moller PL, Kristmundsdottir S, Eggertsson HP, et al. Ratatosk: hybrid error correction of long reads enables accurate variant calling and assembly. Genome Biol. 2021;22(1):28.

10. Wagner J, Olson ND, Harris L, Khan Z, Farek J, Mahmoud M, et al. Benchmarking challenging small variants with linked and long reads. Cell Genom. 2022;2(5).

11. Shafin K, Pesout T, Lorig-Roach R, Haukness M, Olsen HE, Bosworth C, et al. Nanopore sequencing and the Shasta toolkit enable efficient de novo assembly of eleven human genomes. Nat Biotechnol. 2020;38(9):1044–53.

12. Li H. Minimap2: pairwise alignment for nucleotide sequences. Bioinformatics. 2018;34(18):3094–100.

13. Li H, Durbin R. Fast and accurate short read alignment with Burrows-Wheeler transform. Bioinformatics. 2009;25(14):1754–60.

14. Danecek P, Bonfield JK, Liddle J, Marshall J, Ohan V, Pollard MO, et al. Twelve years of SAMtools and BCFtools. Gigascience. 2021;10(2).

15. Krusche P, Trigg L, Boutros PC, Mason CE, De La Vega FM, Moore BL, et al. Best practices for benchmarking germline small-variant calls in human genomes. Nat Biotechnol. 2019;37(5):555–60.

16. Robinson JT, Thorvaldsdottir H, Winckler W, Guttman M, Lander ES, Getz G, et al. Integrative genomics viewer. Nat Biotechnol. 2011;29(1):24–6.

17. Aganezov S, Yan SM, Soto DC, Kirsche M, Zarate S, Avdeyev P, et al. A complete reference genome improves analysis of human genetic variation. Science. 2022;376(6588):eabl3533.

18. Prodanov T, Bansal V. Sensitive alignment using paralogous sequence variants improves long-read mapping and variant calling in segmental duplications. Nucleic Acids Res. 2020;48(19):e114.

19. Patterson M, Marschall T, Pisanti N, van Iersel L, Stougie L, Klau GW, et al. WhatsHap: Weighted Haplotype Assembly for Future-Generation Sequencing Reads. J Comput Biol. 2015;22(6):498–509.

20. Su J, Zheng Z, Ahmed SS, Lam TW, Luo R. Clair3-trio: high-performance Nanopore long-read variant calling in family trios with trio-to-trio deep neural networks. Brief Bioinform. 2022;23(5).

