## Supplementary Materials for "Boosting variant-calling performance with multi-platform sequencing data using Clair3-MP"

### Supplementary Figures

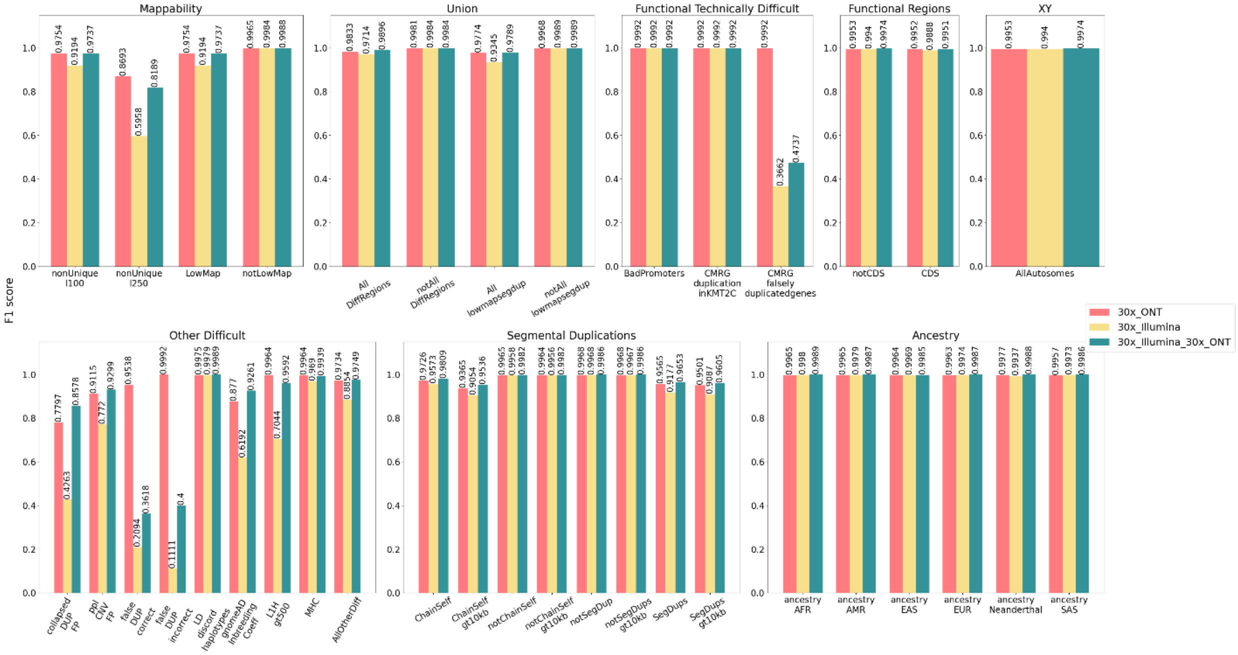

Figure S1. The F1 score of SNP-calling using 30x single-platform vs 30x ONT-Illumina, in eight major stratifications.

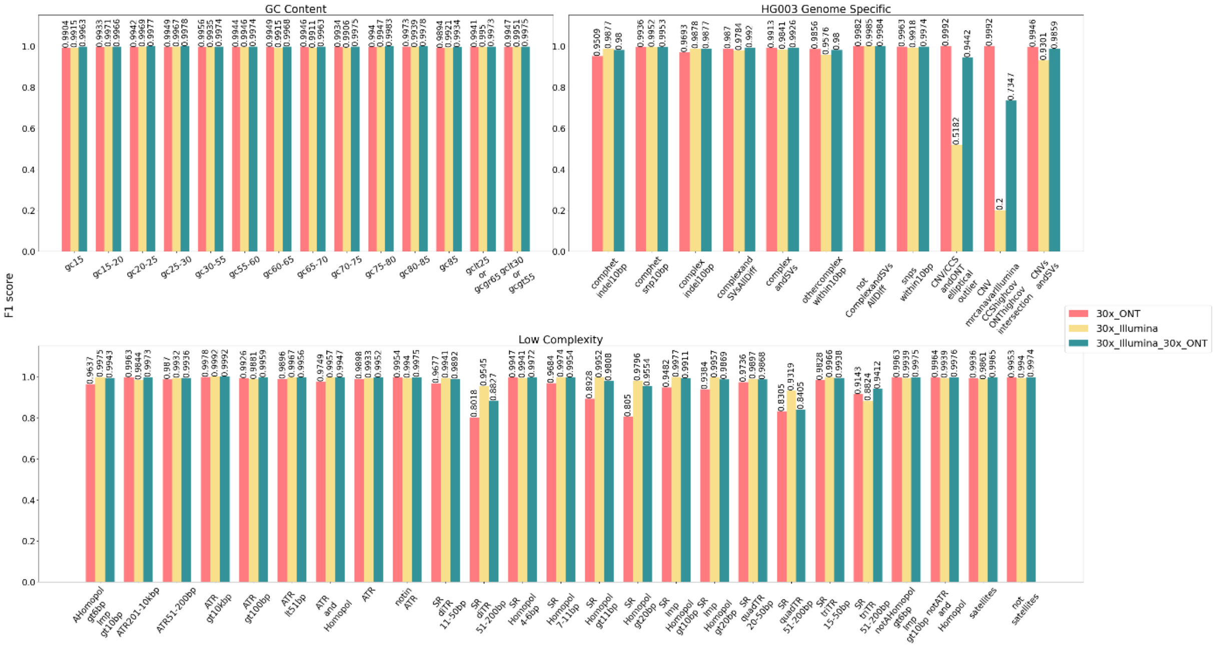

Figure S2. The F1 score of SNP-calling using 30x single-platform vs 30x ONT-Illumina, in three major stratifications.

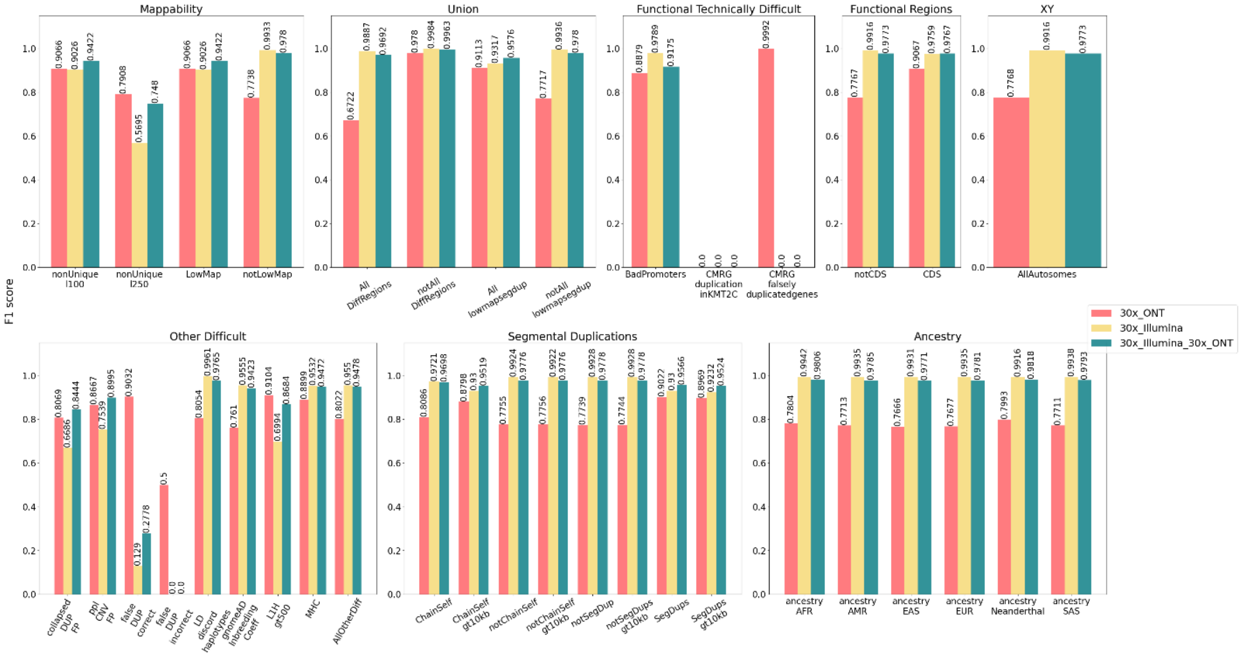

Figure S3. The F1 score of Indel-calling using 30x single-platform vs 30x ONT-Illumina, in eight major stratifications.

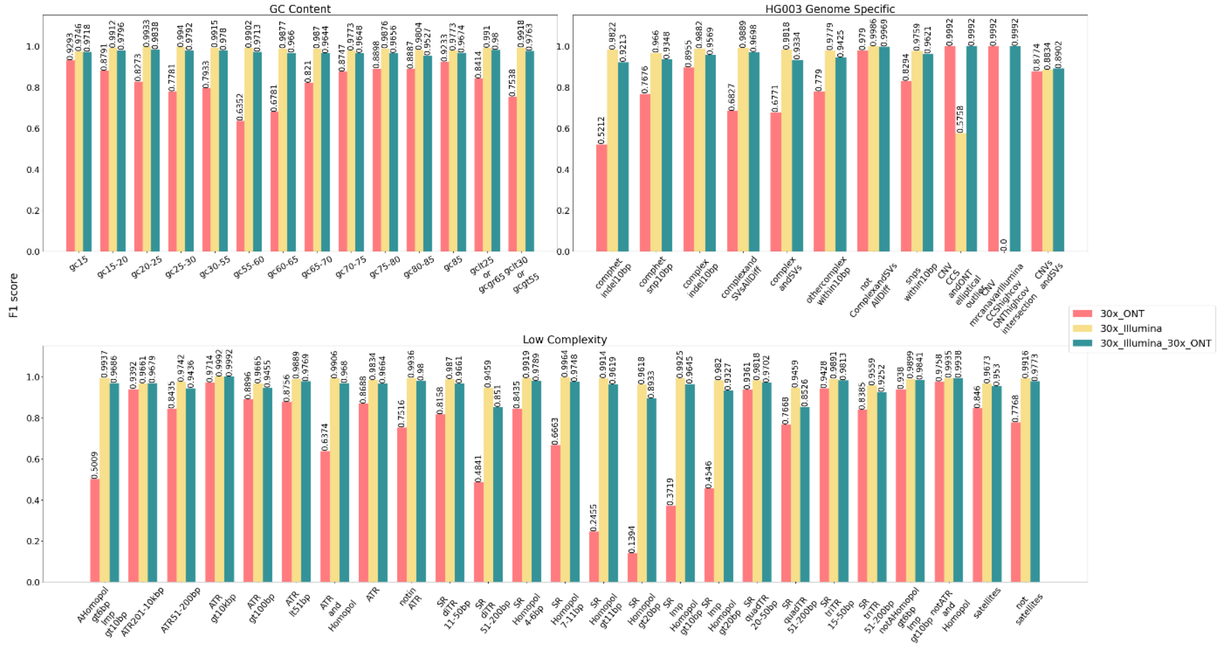

Figure S4. The F1 score of Indel-calling using 30x single-platform vs 30x ONT-Illumina, in three major stratifications.

A.

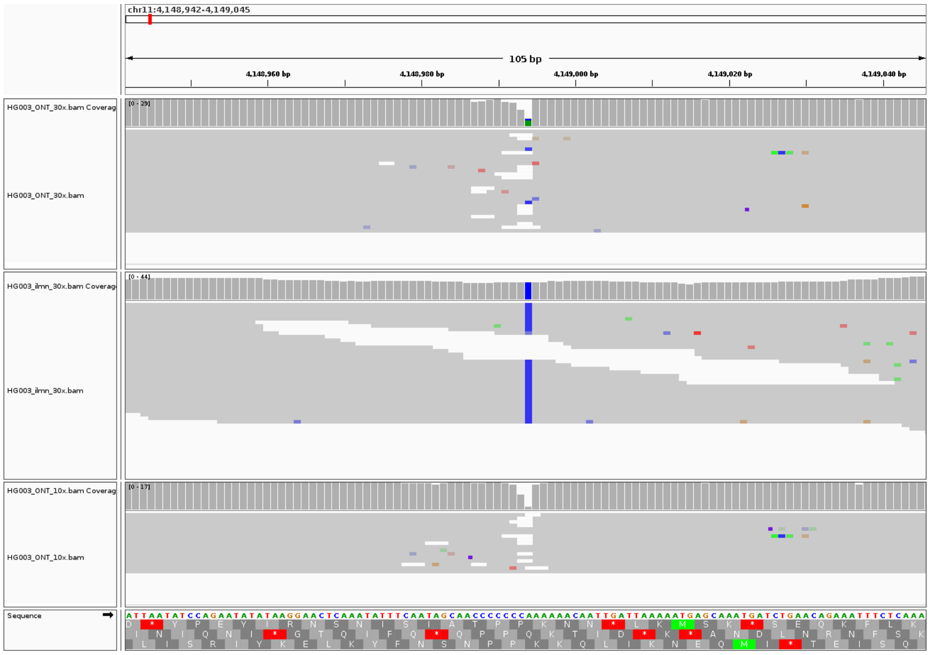

B.

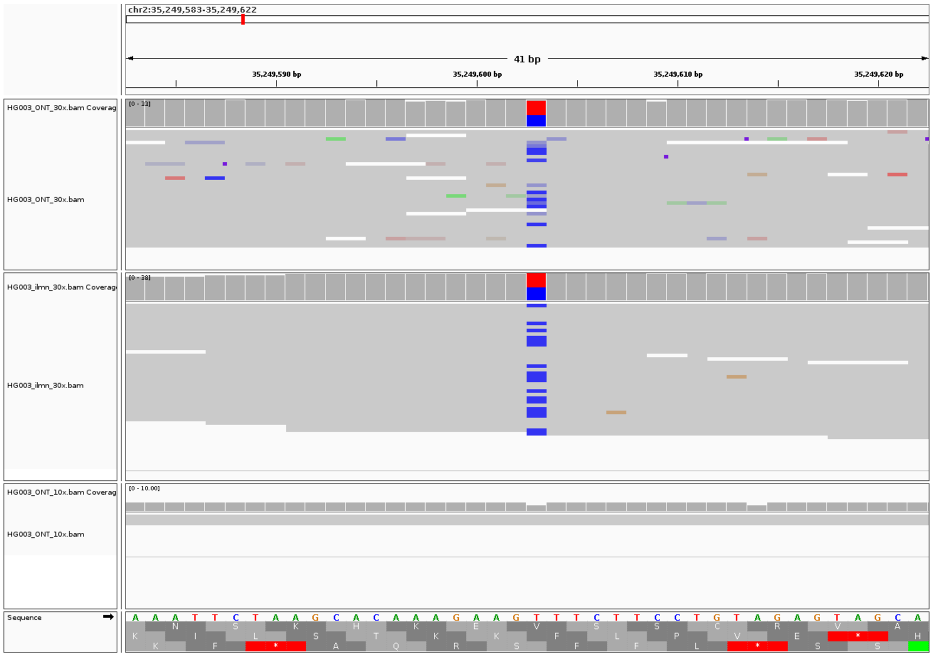

#### Figure S5. Two true SNP cases that were not called by Clair3-MP with 10x ONT-30x Illumina.

A) The variant lacks support from ONT reads. B) The genomic region that the variant locates lacks ONT reads. These cases are both called by Clair3-MP with 30x ONT-Illumina and Clair3 with 30x Illumina.

A.

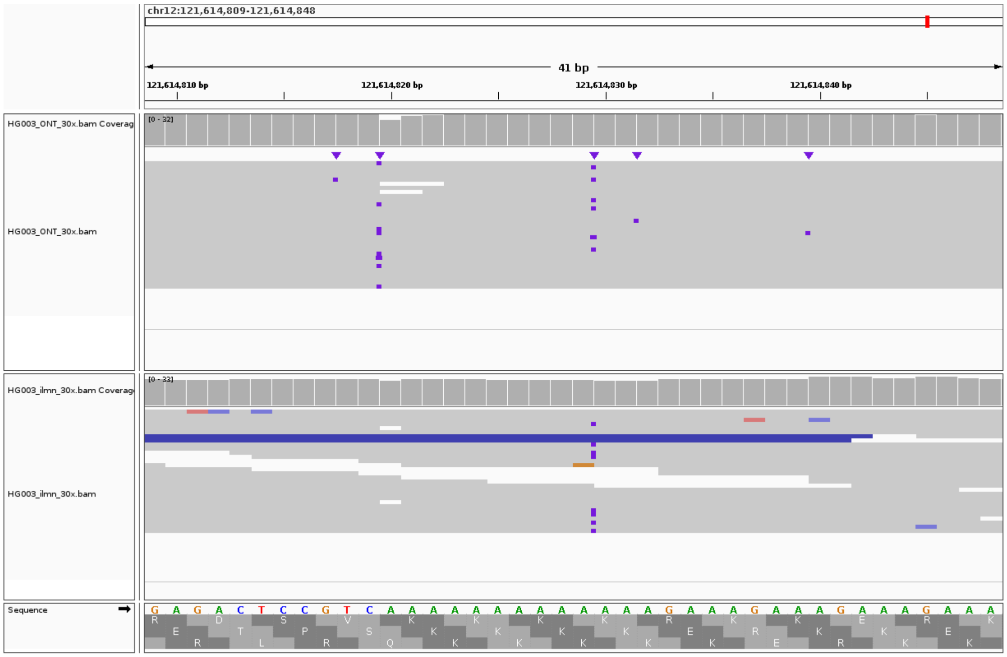

B.

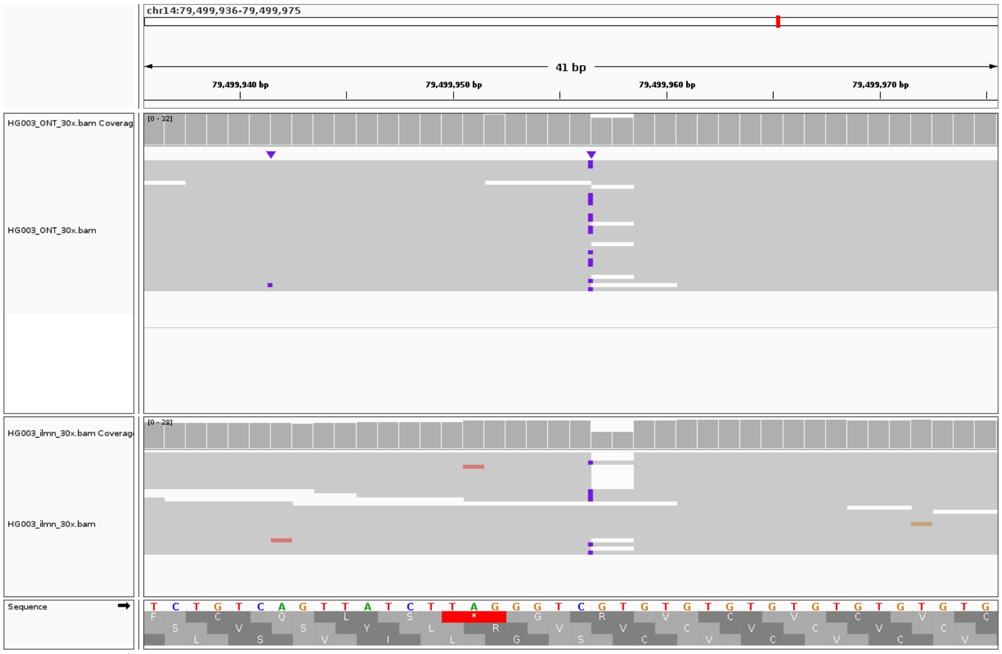

#### Figure S6. Two true Indel cases that were not called by Clair3-MP with 30x ONT-Illumina.

AB) Both have Illumina reads supporting the true allele with the correct insertion sequence. The insertion sequences supported by ONT reads are different in each ONT read.

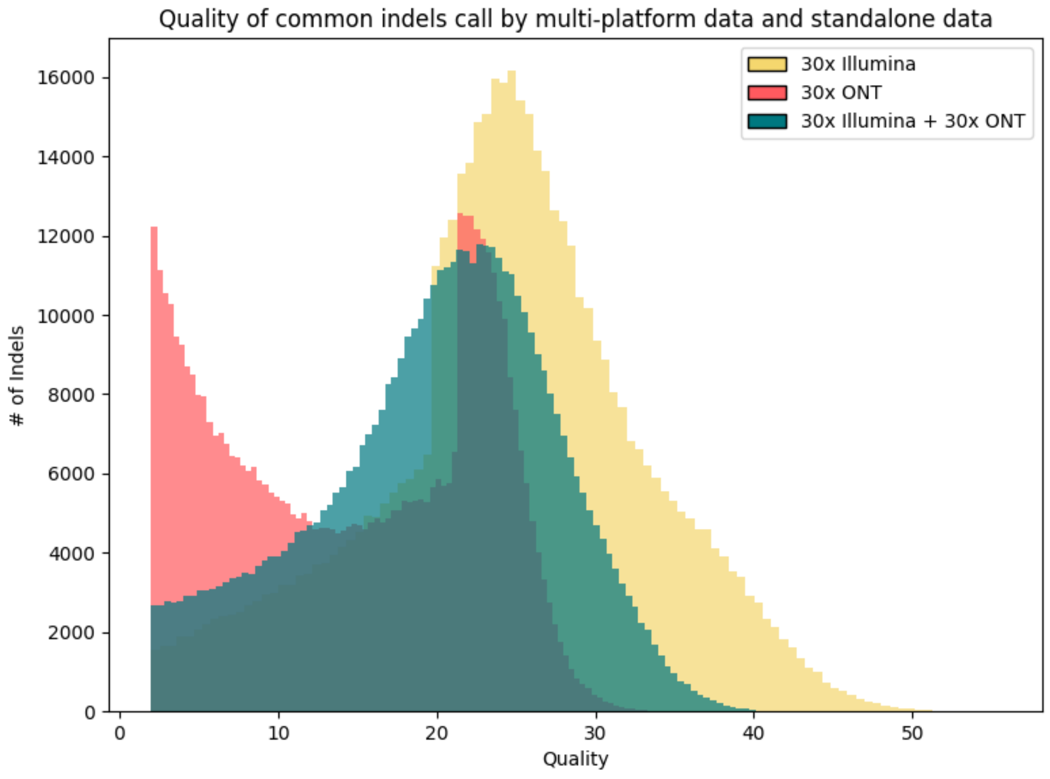

Figure S7. A histogram of the quality scores of the overlapped Indels.

These indels are called by Clair3 with 30x Illumina, Clair3 with 30x ONT, and Clair3-MP with ONT-Illumina (both in 30x) datasets. Despite these indels can be called in the multi-platform data, the quality score of them has a decreasing trend compared to that being called by the Illumina data.

### Supplementary Tables

#### Table S1. The precision, recall and F1 score of variant calling using 30x single- vs 30x multi-platform data (ONT and Illumina, PacBio and Illumina, PacBio, and ONT), based on the variant types.

| | Coverages | | | Overall | | | SNPs | | | Indels | | |
|---|---|---|---|---|---|---|---|---|---|---|---|---|
| Software | ON | Illuminaa | PacBio | Precisionn | Recall | F1 score | Precision | Recall | F1 score | Precision | Recall | F1 score |
| Clair3 | 10x | / | / | 0.9605 | 0.9016 | 0.9301 | 0.9715 | 0.9648 | 0.9681 | 0.8386 | 0.4843 | 0.6140 |
| | 20x | / | / | 0.9816 | 0.9453 | 0.9631 | 0.9921 | 0.9943 | 0.9932 | 0.8847 | 0.6221 | 0.7305 |
| | 30x | / | / | 0.9857 | 0.9541 | 0.9696 | 0.9950 | 0.9956 | 0.9953 | 0.9055 | 0.6803 | 0.7769 |
| | / | 10x | / | 0.9240 | 0.9432 | 0.9335 | 0.9226 | 0.9485 | 0.9353 | 0.9333 | 0.9081 | 0.9205 |
| | / | 20x | / | 0.9893 | 0.9867 | 0.9880 | 0.9899 | 0.9882 | 0.9890 | 0.9857 | 0.9765 | 0.9811 |
| | / | 30x | / | 0.9950 | 0.9923 | 0.9937 | 0.9951 | 0.9928 | 0.9940 | 0.9943 | 0.9890 | 0.9916 |
| | / | / | 10x | 0.9816 | 0.9801 | 0.9809 | 0.9922 | 0.9876 | 0.9899 | 0.9156 | 0.9308 | 0.9231 |
| | / | / | 20x | 0.9965 | 0.9969 | 0.9967 | 0.9986 | 0.9986 | 0.9986 | 0.9828 | 0.9854 | 0.9841 |
| | / | / | 30x | 0.9982 | 0.9983 | 0.9983 | 0.9990 | 0.9990 | 0.9990 | 0.9928 | 0.9938 | **0.9933** |
| Clair3-MP (ONT-Illumina) | 10x | 10x | / | 0.9822 | 0.9609 | 0.9714 | 0.9961 | 0.9794 | 0.9877 | 0.8891 | 0.8388 | 0.8632 |
| | 10x | 20x | / | 0.9852 | 0.9725 | 0.9788 | 0.9974 | 0.9871 | 0.9922 | 0.9058 | 0.8767 | 0.8910 |
| | 10x | 30x | / | 0.9860 | 0.9739 | 0.9799 | 0.9978 | 0.9877 | 0.9927 | 0.9093 | 0.8830 | 0.8959 |
| | 20x | 10x | / | 0.9910 | 0.9800 | 0.9855 | 0.9979 | 0.9902 | 0.9940 | 0.9462 | 0.9131 | 0.9293 |
| | 20x | 20x | / | 0.9938 | 0.9894 | 0.9916 | 0.9983 | 0.9954 | 0.9968 | 0.9651 | 0.9503 | 0.9577 |
| | 20x | 30x | / | 0.9946 | 0.9906 | 0.9926 | 0.9985 | 0.9957 | 0.9971 | 0.9694 | 0.9570 | 0.9631 |
| | 30x | 10x | / | 0.9931 | 0.9830 | 0.9880 | 0.9983 | 0.9913 | 0.9948 | 0.9595 | 0.9285 | 0.9438 |
| | 30x | 20x | / | 0.9959 | 0.9917 | 0.9938 | 0.9985 | 0.9957 | 0.9971 | 0.9788 | 0.9653 | 0.9720 |
| | 30x | 30x | / | 0.9966 | 0.9929 | 0.9947 | 0.9987 | 0.9961 | 0.9974 | 0.9831 | 0.9717 | 0.9774 |
| Clair3-MP (PacBio- | / | 10x | 10x | 0.9960 | 0.9873 | 0.9916 | 0.9985 | 0.9906 | 0.9945 | 0.9802 | 0.9653 | 0.9727 |
| | / | 20x | 20x | 0.9988 | 0.9967 | 0.9978 | 0.9994 | 0.9971 | 0.9982 | 0.9954 | 0.9942 | 0.9948 |
| | / | 30x | 30x | 0.9992 | 0.9974 | 0.9983 | 0.9995 | 0.9976 | 0.9985 | 0.9973 | 0.9966 | **0.9969** |
| Clair3-MP (ONT- | 10x | / | 10x | 0.9834 | 0.9664 | 0.9748 | 0.9974 | 0.9856 | 0.9914 | 0.8894 | 0.8403 | 0.8642 |
| | 20x | / | 20x | 0.9938 | 0.9918 | 0.9928 | 0.9987 | 0.9979 | 0.9983 | 0.9626 | 0.9515 | 0.9570 |
| | 30x | / | 30x | 0.9965 | 0.9950 | 0.9957 | 0.9988 | 0.9983 | 0.9986 | 0.9812 | 0.9731 | 0.9771 |

Table S2. The precision, recall, and F1 score of variant calling using 30x ONT data, stratifying into all defined genomic regions and based on the variant types.

| | | SNPs | | | Indels | | |
|---|---|---|---|---|---|---|---|
| Stratification Type | Stratification Group | Precision | Recall | F1 score | Precision | Recall | F1 score |
| LowComplexity | AHomopol_gt6bp_Imp_gt10bp | 0.9599 | 0.9675 | 0.9637 | 0.7859 | 0.3676 | 0.5009 |
| | ATR201-10kbp | 0.9967 | 0.9959 | 0.9963 | 0.9569 | 0.9222 | 0.9392 |
| | ATR51-200bp | 0.9849 | 0.9891 | 0.9870 | 0.8715 | 0.8173 | 0.8435 |
| | ATR_gt10kbp | 0.9992 | 0.9956 | 0.9978 | 0.9992 | 0.9444 | 0.9714 |
| | ATR_gt100bp | 0.9917 | 0.9935 | 0.9926 | 0.9178 | 0.8632 | 0.8896 |
| | ATR_lt51bp | 0.9884 | 0.9908 | 0.9896 | 0.8897 | 0.8619 | 0.8756 |
| | ATR_and_Homopol | 0.9722 | 0.9776 | 0.9749 | 0.8360 | 0.5151 | 0.6374 |
| | ATR | 0.9886 | 0.9911 | 0.9898 | 0.8872 | 0.8511 | 0.8688 |
| | notin_ATR | 0.9952 | 0.9957 | 0.9954 | 0.9116 | 0.6394 | 0.7516 |
| | SR_diTR_11-50bp | 0.9626 | 0.9730 | 0.9677 | 0.8417 | 0.7915 | 0.8158 |
| | SR_diTR_51-200bp | 0.7447 | 0.8684 | 0.8018 | 0.5139 | 0.4576 | 0.4841 |
| | SR_diTR_gt200bp | 0.0000 | 0.0000 | None | 0.0000 | 0.0000 | None |
| | SR_Homopol_4-6bp | 0.9946 | 0.9947 | 0.9947 | 0.9270 | 0.7738 | 0.8435 |
| | SR_Homopol_7-11bp | 0.9653 | 0.9715 | 0.9684 | 0.8256 | 0.5586 | 0.6663 |
| | SR_Homopol_gt11bp | 0.8820 | 0.9038 | 0.8928 | 0.6365 | 0.1521 | 0.2455 |
| | SR_Homopol_gt20bp | 0.8017 | 0.8084 | 0.8050 | 0.4890 | 0.0813 | 0.1394 |
| | SR_Imp_Homopol_gt10bp | 0.9428 | 0.9537 | 0.9482 | 0.7263 | 0.2499 | 0.3719 |
| | SR_Imp_Homopol_gt20bp | 0.9327 | 0.9442 | 0.9384 | 0.7844 | 0.3201 | 0.4546 |
| | SR_quadTR_20-50bp | 0.9699 | 0.9773 | 0.9736 | 0.9595 | 0.9139 | 0.9361 |
| | SR_quadTR_51-200bp | 0.8534 | 0.8087 | 0.8305 | 0.8221 | 0.7185 | 0.7668 |
| | SR_quadTR_gt200bp | None | 0.0000 | None | None | 0.0000 | None |
| | SR_triTR_15-50bp | 0.9797 | 0.9859 | 0.9828 | 0.9670 | 0.9198 | 0.9428 |
| | SR_triTR_51-200bp | 0.9412 | 0.8889 | 0.9143 | 0.8650 | 0.8136 | 0.8385 |
| | SR_triTR_gt200bp | None | 0.0000 | None | None | 0.0000 | None |

| | | | | | | | |
|---|---|---|---|---|---|---|---|
| | notAHomopol_gt6bp_Imp_gt10bp | 0.9961 | 0.9964 | 0.9963 | 0.9492 | 0.9271 | 0.9380 |
| | notATR_and_Homopol | 0.9963 | 0.9966 | 0.9964 | 0.9843 | 0.9674 | 0.9758 |
| | satellites | 0.9936 | 0.9936 | 0.9936 | 0.9279 | 0.7773 | 0.8460 |
| | not_satellites | 0.9950 | 0.9956 | 0.9953 | 0.9052 | 0.6803 | 0.7768 |
| SegmentalDuplications | ChainSelf | 0.9586 | 0.9871 | 0.9726 | 0.9094 | 0.7280 | 0.8086 |
| | ChainSelf_gt10kb | 0.9031 | 0.9725 | 0.9365 | 0.8998 | 0.8607 | 0.8798 |
| | gt5SegDups_gt10kb | None | 0.0000 | None | None | 0.0000 | None |
| | notChainSelf | 0.9969 | 0.9960 | 0.9965 | 0.9050 | 0.6784 | 0.7755 |
| | notChainSelf_gt10kb | 0.9968 | 0.9960 | 0.9964 | 0.9052 | 0.6785 | 0.7756 |
| | notSegDup | 0.9975 | 0.9962 | 0.9968 | 0.9049 | 0.6761 | 0.7739 |
| | notSegDups_gt10kb | 0.9975 | 0.9962 | 0.9968 | 0.9051 | 0.6768 | 0.7744 |
| | SegDups | 0.9339 | 0.9803 | 0.9565 | 0.9173 | 0.8877 | 0.9023 |
| | SegDups_gt10kb | 0.9242 | 0.9776 | 0.9501 | 0.9100 | 0.8842 | 0.8969 |
| Mappability | nonUnique_l100 | 0.9650 | 0.9862 | 0.9754 | 0.9291 | 0.8851 | 0.9066 |
| | nonUnique_l250 | 0.8340 | 0.9076 | 0.8693 | 0.8143 | 0.7685 | 0.7908 |
| | LowMap | 0.9650 | 0.9862 | 0.9754 | 0.9291 | 0.8851 | 0.9066 |
| | notLowMap | 0.9969 | 0.9962 | 0.9965 | 0.9045 | 0.6760 | 0.7738 |
| OtherDiffcult | KIR | None | 0.0000 | None | None | 0.0000 | None |
| | collapsed_DUP_FP | 0.6543 | 0.9646 | 0.7797 | 0.7520 | 0.8704 | 0.8069 |
| | ppl_CNV_FP | 0.8703 | 0.9567 | 0.9115 | 0.8783 | 0.8555 | 0.8667 |
| | false_DUP_correct | 0.9355 | 0.9727 | 0.9538 | 0.8485 | 0.9655 | 0.9032 |
| | false_DUP_incorrect | 0.9992 | 0.9992 | 0.9992 | 0.5000 | 0.5000 | 0.5000 |
| | LD_discord_haplotypes | 0.9976 | 0.9974 | 0.9975 | 0.9157 | 0.7188 | 0.8054 |
| | gnomeAD_InbreedingCoeff | 0.8271 | 0.9333 | 0.8770 | 0.8851 | 0.6674 | 0.7610 |
| | L1H_gt500 | 0.9981 | 0.9946 | 0.9964 | 0.9681 | 0.8592 | 0.9104 |
| | MHC | 0.9977 | 0.9952 | 0.9964 | 0.9243 | 0.8580 | 0.8899 |
| | VDJ | None | 0.0000 | None | None | 0.0000 | None |
| | contigs_lt500kb | None | 0.0000 | None | None | 0.0000 | None |

| | | | | | | | |
|---|---|---|---|---|---|---|---|
| | gaps_slop15kb | None | 0.0000 | None | None | 0.0000 | None |
| | AllOtherDiff | 0.9617 | 0.9854 | 0.9734 | 0.9011 | 0.7228 | 0.8022 |
| FunctionalRegions | notCDS | 0.9950 | 0.9956 | 0.9953 | 0.9052 | 0.6801 | 0.7767 |
| | CDS | 0.9957 | 0.9948 | 0.9952 | 0.9026 | 0.9108 | 0.9067 |
| FunctionalTechnicallyDifficult Regions | BadPromoters | 0.9992 | 0.9992 | 0.9992 | 0.9531 | 0.8310 | 0.8879 |
| | CMRG_duplicationinKMT2C | 0.9992 | 0.9992 | 0.9992 | None | 0.0000 | None |
| | CMRG_falselyduplicatedgenes | 0.9992 | 0.9992 | 0.9992 | 0.9992 | 0.9992 | 0.9992 |
| GCContent | gc15 | 0.9864 | 0.9945 | 0.9904 | 0.9558 | 0.9042 | 0.9293 |
| | gc15-20 | 0.9933 | 0.9933 | 0.9933 | 0.9485 | 0.8191 | 0.8791 |
| | gc20-25 | 0.9942 | 0.9942 | 0.9942 | 0.9296 | 0.7454 | 0.8273 |
| | gc25-30 | 0.9948 | 0.9949 | 0.9949 | 0.9069 | 0.6813 | 0.7781 |
| | gc30-55 | 0.9954 | 0.9958 | 0.9956 | 0.9053 | 0.7060 | 0.7933 |
| | gc55-60 | 0.9937 | 0.9952 | 0.9944 | 0.8690 | 0.5005 | 0.6352 |
| | gc60-65 | 0.9943 | 0.9954 | 0.9949 | 0.8857 | 0.5493 | 0.6781 |
| | gc65-70 | 0.9934 | 0.9957 | 0.9946 | 0.9282 | 0.7360 | 0.8210 |
| | gc70-75 | 0.9913 | 0.9954 | 0.9934 | 0.9340 | 0.8224 | 0.8747 |
| | gc75-80 | 0.9935 | 0.9945 | 0.9940 | 0.9444 | 0.8412 | 0.8898 |
| | gc80-85 | 0.9990 | 0.9956 | 0.9973 | 0.9613 | 0.8263 | 0.8887 |
| | gc85 | 0.9920 | 0.9868 | 0.9894 | 0.9589 | 0.8903 | 0.9233 |
| | gclt25orgcgr65 | 0.9937 | 0.9946 | 0.9941 | 0.9341 | 0.7655 | 0.8414 |
| | gclt30orgcgt55 | 0.9943 | 0.9951 | 0.9947 | 0.9051 | 0.6459 | 0.7538 |
| union | AllDiffRegions | 0.9784 | 0.9883 | 0.9833 | 0.8536 | 0.5544 | 0.6722 |
| | notAllDiffRegions | 0.9989 | 0.9973 | 0.9981 | 0.9868 | 0.9695 | 0.9780 |
| | Alllowmapsegdup | 0.9666 | 0.9886 | 0.9774 | 0.9334 | 0.8902 | 0.9113 |
| | notAlllowmapsegdup | 0.9975 | 0.9962 | 0.9968 | 0.9040 | 0.6732 | 0.7717 |
| GenomeSpecific_HG003 | comphetindel10bp | 0.9248 | 0.9784 | 0.9509 | 0.7278 | 0.4059 | 0.5212 |
| | comphetsnp10bp | 0.9959 | 0.9913 | 0.9936 | 0.8326 | 0.7121 | 0.7676 |
| | complexindel10bp | 0.9639 | 0.9748 | 0.9693 | 0.9596 | 0.8394 | 0.8955 |

| | | | | | | | |
|---|---|---|---|---|---|---|---|
| | complexandSVs_AllDiff | 0.9837 | 0.9903 | 0.9870 | 0.8591 | 0.5665 | 0.6827 |
| | complexandSVs | 0.9922 | 0.9903 | 0.9913 | 0.8200 | 0.5767 | 0.6771 |
| | othercomplexwithin10bp | 0.9877 | 0.9836 | 0.9856 | 0.9064 | 0.6830 | 0.7790 |
| | notComplexandSVs_AllDiff | 0.9989 | 0.9974 | 0.9982 | 0.9873 | 0.9708 | 0.9790 |
| | snpswithin10bp | 0.9985 | 0.9942 | 0.9963 | 0.8795 | 0.7847 | 0.8294 |
| | CNV_CCSandONT_elliptical_outlier | 0.9992 | 0.9992 | 0.9992 | 0.9992 | 0.9992 | 0.9992 |
| | CNV_mrcanavarIllumina_CCShighcov_ ONThighcov_intersection | 0.9992 | 0.9992 | 0.9992 | 0.9992 | 0.9992 | 0.9992 |
| | SV_pbsv_slop25percent | None | 0.0000 | None | None | 0.0000 | None |
| | CNVsandSVs | 0.9951 | 0.9941 | 0.9946 | 0.9140 | 0.8436 | 0.8774 |
| XY | chrX_PAR | 0.0000 | 0.0000 | None | 0.0000 | 0.0000 | None |
| | chrX_XTR | 0.0000 | 0.0000 | None | 0.0000 | 0.0000 | None |
| | chrX_ampliconic | 0.0000 | 0.0000 | None | 0.0000 | 0.0000 | None |
| | chrX_nonPAR | 0.0000 | 0.0000 | None | 0.0000 | 0.0000 | None |
| | chrY_PAR | 0.0000 | 0.0000 | None | 0.0000 | 0.0000 | None |
| | chrY_XTR | 0.0000 | 0.0000 | None | 0.0000 | 0.0000 | None |
| | chrY_ampliconic | 0.0000 | 0.0000 | None | 0.0000 | 0.0000 | None |
| | chrY_nonPAR | 0.0000 | 0.0000 | None | 0.0000 | 0.0000 | None |
| | AllAutosomes | 0.9950 | 0.9956 | 0.9953 | 0.9052 | 0.6803 | 0.7768 |
| Ancestry | ancestry_AFR | 0.9964 | 0.9965 | 0.9965 | 0.9107 | 0.6827 | 0.7804 |
| | ancestry_AMR | 0.9968 | 0.9961 | 0.9965 | 0.9027 | 0.6733 | 0.7713 |
| | ancestry_EAS | 0.9966 | 0.9962 | 0.9964 | 0.8996 | 0.6679 | 0.7666 |
| | ancestry_EUR | 0.9968 | 0.9959 | 0.9963 | 0.8996 | 0.6695 | 0.7677 |
| | ancestry_Neanderthal | 0.9982 | 0.9972 | 0.9977 | 0.9180 | 0.7078 | 0.7993 |
| | ancestry_SAS | 0.9951 | 0.9963 | 0.9957 | 0.9036 | 0.6725 | 0.7711 |

Table S3. The precision, recall, and F1 score of variant calling using 30x Illumina data, stratifying into all defined genomic regions and based on the variant types

| | | SNPs | | | Indels | | |
|---|---|---|---|---|---|---|---|
| Stratification Type | Stratification Group | Precision | Recall | F1 score | Precision | Recall | F1 score |
| LowComplexity | AHomopol_gt6bp_Imp_gt10bp | 0.9973 | 0.9977 | 0.9975 | 0.9949 | 0.9925 | 0.9937 |
| | ATR201-10kbp | 0.9814 | 0.9873 | 0.9844 | 0.9812 | 0.9515 | 0.9661 |
| | ATR51-200bp | 0.9913 | 0.9952 | 0.9932 | 0.9846 | 0.9640 | 0.9742 |
| | ATR_gt10kbp | 0.9992 | 0.9992 | 0.9992 | 0.9992 | 0.9992 | 0.9992 |
| | ATR_gt100bp | 0.9854 | 0.9909 | 0.9881 | 0.9806 | 0.9529 | 0.9665 |
| | ATR_lt51bp | 0.9961 | 0.9973 | 0.9967 | 0.9929 | 0.9849 | 0.9889 |
| | ATR_and_Homopol | 0.9950 | 0.9964 | 0.9957 | 0.9933 | 0.9878 | 0.9906 |
| | ATR | 0.9919 | 0.9948 | 0.9933 | 0.9899 | 0.9770 | 0.9834 |
| | notin_ATR | 0.9952 | 0.9928 | 0.9940 | 0.9954 | 0.9919 | 0.9936 |
| | SR_diTR_11-50bp | 0.9928 | 0.9953 | 0.9941 | 0.9909 | 0.9832 | 0.9870 |
| | SR_diTR_51-200bp | 0.9130 | 0.9992 | 0.9545 | 0.9662 | 0.9265 | 0.9459 |
| | SR_diTR_gt200bp | 0.0000 | 0.0000 | None | 0.0000 | 0.0000 | None |
| | SR_Homopol_4-6bp | 0.9952 | 0.9931 | 0.9941 | 0.9948 | 0.9891 | 0.9919 |
| | SR_Homopol_7-11bp | 0.9973 | 0.9975 | 0.9974 | 0.9974 | 0.9954 | 0.9964 |
| | SR_Homopol_gt11bp | 0.9946 | 0.9957 | 0.9952 | 0.9926 | 0.9903 | 0.9914 |
| | SR_Homopol_gt20bp | 0.9800 | 0.9791 | 0.9796 | 0.9699 | 0.9537 | 0.9618 |
| | SR_Imp_Homopol_gt10bp | 0.9974 | 0.9979 | 0.9977 | 0.9938 | 0.9912 | 0.9925 |
| | SR_Imp_Homopol_gt20bp | 0.9959 | 0.9956 | 0.9957 | 0.9863 | 0.9778 | 0.9820 |
| | SR_quadTR_20-50bp | 0.9876 | 0.9918 | 0.9897 | 0.9896 | 0.9741 | 0.9818 |
| | SR_quadTR_51-200bp | 0.9333 | 0.9304 | 0.9319 | 0.9719 | 0.9212 | 0.9459 |
| | SR_quadTR_gt200bp | None | 0.0000 | None | None | 0.0000 | None |
| | SR_triTR_15-50bp | 0.9955 | 0.9976 | 0.9966 | 0.9936 | 0.9847 | 0.9891 |
| | SR_triTR_51-200bp | 0.9375 | 0.8333 | 0.8824 | 0.9747 | 0.9379 | 0.9559 |
| | SR_triTR_gt200bp | None | 0.0000 | None | None | 0.0000 | None |

| | | | | | | | |
|---|---|---|---|---|---|---|---|
| | notAHomopol_gt6bp_Imp_gt10bp | 0.9951 | 0.9927 | 0.9939 | 0.9937 | 0.9862 | 0.9899 |
| | notATR_and_Homopol | 0.9952 | 0.9926 | 0.9939 | 0.9960 | 0.9910 | 0.9935 |
| | satellites | 0.9866 | 0.9856 | 0.9861 | 0.9795 | 0.9555 | 0.9673 |
| | not_satellites | 0.9952 | 0.9928 | 0.9940 | 0.9943 | 0.9890 | 0.9916 |
| SegmentalDuplications | ChainSelf | 0.9608 | 0.9539 | 0.9573 | 0.9803 | 0.9641 | 0.9721 |
| | ChainSelf_gt10kb | 0.9183 | 0.8929 | 0.9054 | 0.9542 | 0.9069 | 0.9300 |
| | gt5SegDups_gt10kb | None | 0.0000 | None | None | 0.0000 | None |
| | notChainSelf | 0.9969 | 0.9948 | 0.9958 | 0.9948 | 0.9900 | 0.9924 |
| | notChainSelf_gt10kb | 0.9965 | 0.9946 | 0.9956 | 0.9946 | 0.9898 | 0.9922 |
| | notSegDup | 0.9976 | 0.9961 | 0.9968 | 0.9950 | 0.9907 | 0.9928 |
| | notSegDups_gt10kb | 0.9975 | 0.9959 | 0.9967 | 0.9950 | 0.9906 | 0.9928 |
| | SegDups | 0.9284 | 0.9073 | 0.9177 | 0.9538 | 0.9073 | 0.9300 |
| | SegDups_gt10kb | 0.9207 | 0.8970 | 0.9087 | 0.9492 | 0.8987 | 0.9232 |
| Mappability | nonUnique_l100 | 0.9462 | 0.8940 | 0.9194 | 0.9515 | 0.8585 | 0.9026 |
| | nonUnique_l250 | 0.7370 | 0.5000 | 0.5958 | 0.7494 | 0.4593 | 0.5695 |
| | LowMap | 0.9462 | 0.8940 | 0.9194 | 0.9515 | 0.8585 | 0.9026 |
| | notLowMap | 0.9980 | 0.9989 | 0.9984 | 0.9950 | 0.9916 | 0.9933 |
| OtherDiffcult | KIR | None | 0.0000 | None | None | 0.0000 | None |
| | collapsed_DUP_FP | 0.2846 | 0.8493 | 0.4263 | 0.5503 | 0.8519 | 0.6686 |
| | ppl_CNV_FP | 0.7450 | 0.8010 | 0.7720 | 0.7722 | 0.7365 | 0.7539 |
| | false_DUP_correct | 0.7532 | 0.1216 | 0.2094 | 0.9992 | 0.0690 | 0.1290 |
| | false_DUP_incorrect | 0.5000 | 0.0625 | 0.1111 | None | 0.0000 | None |
| | LD_discord_haplotypes | 0.9983 | 0.9975 | 0.9979 | 0.9980 | 0.9942 | 0.9961 |
| | gnomeAD_InbreedingCoeff | 0.5022 | 0.8075 | 0.6192 | 0.9445 | 0.9668 | 0.9555 |
| | L1H_gt500 | 0.8694 | 0.5921 | 0.7044 | 0.9098 | 0.5681 | 0.6994 |
| | MHC | 0.9572 | 0.9810 | 0.9690 | 0.9514 | 0.9550 | 0.9532 |
| | VDJ | None | 0.0000 | None | None | 0.0000 | None |
| | contigs_lt500kb | None | 0.0000 | None | None | 0.0000 | None |

| | | | | | | | |
|---|---|---|---|---|---|---|---|
| | gaps_slop15kb | None | 0.0000 | None | None | 0.0000 | None |
| | AllOtherDiff | 0.8662 | 0.9055 | 0.8854 | 0.9562 | 0.9539 | 0.9550 |
| FunctionalRegions | notCDS | 0.9952 | 0.9928 | 0.9940 | 0.9943 | 0.9890 | 0.9916 |
| | CDS | 0.9881 | 0.9894 | 0.9888 | 0.9761 | 0.9757 | 0.9759 |
| FunctionalTechnicallyDifficult Regions | BadPromoters | 0.9992 | 0.9992 | 0.9992 | 0.9861 | 0.9718 | 0.9789 |
| | CMRG_duplicationinKMT2C | 0.9992 | 0.9992 | 0.9992 | None | 0.0000 | None |
| | CMRG_falselyduplicatedgenes | 0.9992 | 0.2241 | 0.3662 | None | 0.0000 | None |
| GCContent | gc15 | 0.9879 | 0.9951 | 0.9915 | 0.9811 | 0.9682 | 0.9746 |
| | gc15-20 | 0.9975 | 0.9968 | 0.9971 | 0.9945 | 0.9880 | 0.9912 |
| | gc20-25 | 0.9967 | 0.9970 | 0.9969 | 0.9951 | 0.9916 | 0.9933 |
| | gc25-30 | 0.9968 | 0.9966 | 0.9967 | 0.9955 | 0.9925 | 0.9940 |
| | gc30-55 | 0.9951 | 0.9919 | 0.9935 | 0.9944 | 0.9885 | 0.9915 |
| | gc55-60 | 0.9949 | 0.9942 | 0.9946 | 0.9926 | 0.9878 | 0.9902 |
| | gc60-65 | 0.9930 | 0.9900 | 0.9915 | 0.9919 | 0.9836 | 0.9877 |
| | gc65-70 | 0.9916 | 0.9905 | 0.9911 | 0.9913 | 0.9827 | 0.9870 |
| | gc70-75 | 0.9884 | 0.9928 | 0.9906 | 0.9843 | 0.9704 | 0.9773 |
| | gc75-80 | 0.9937 | 0.9958 | 0.9947 | 0.9876 | 0.9876 | 0.9876 |
| | gc80-85 | 0.9907 | 0.9970 | 0.9939 | 0.9845 | 0.9763 | 0.9804 |
| | gc85 | 0.9947 | 0.9894 | 0.9921 | 0.9805 | 0.9742 | 0.9773 |
| | gclt25orgcgr65 | 0.9948 | 0.9952 | 0.9950 | 0.9935 | 0.9884 | 0.9910 |
| | gclt30orgcgt55 | 0.9954 | 0.9948 | 0.9951 | 0.9940 | 0.9896 | 0.9918 |
| union | AllDiffRegions | 0.9781 | 0.9648 | 0.9714 | 0.9924 | 0.9851 | 0.9887 |
| | notAllDiffRegions | 0.9991 | 0.9985 | 0.9984 | 0.9988 | 0.9980 | 0.9984 |
| | Alllowmapsegdup | 0.9527 | 0.9170 | 0.9345 | 0.9630 | 0.9023 | 0.9317 |
| | notAlllowmapsegdup | 0.9986 | 0.9984 | 0.9989 | 0.9952 | 0.9919 | 0.9936 |
| GenomeSpecific_HG003 | comphetindel10bp | 0.9835 | 0.9919 | 0.9877 | 0.9870 | 0.9775 | 0.9822 |
| | comphetsnp10bp | 0.9972 | 0.9932 | 0.9952 | 0.9631 | 0.9689 | 0.9660 |
| | complexindel10bp | 0.9938 | 0.9819 | 0.9878 | 0.9951 | 0.9813 | 0.9882 |

| | | | | | | | |
|---|---|---|---|---|---|---|---|
| | complexandSVs_AllDiff | 0.9837 | 0.9732 | 0.9784 | 0.9925 | 0.9853 | 0.9889 |
| | complexandSVs | 0.9950 | 0.9734 | 0.9841 | 0.9889 | 0.9749 | 0.9818 |
| | othercomplexwithin10bp | 0.9922 | 0.9254 | 0.9576 | 0.9917 | 0.9644 | 0.9779 |
| | notComplexandSVs_AllDiff | 0.9990 | 0.9987 | 0.9985 | 0.9989 | 0.9984 | 0.9986 |
| | snpswithin10bp | 0.9977 | 0.9860 | 0.9918 | 0.9866 | 0.9653 | 0.9759 |
| | CNV_CCSandONT_elliptical_outlier | 0.3699 | 0.8652 | 0.5182 | 0.4130 | 0.9500 | 0.5758 |
| | CNV_mrcanavarIllumina_CCShighcov_ ONThighcov_intersection | 0.1204 | 0.5909 | 0.2000 | 0.0000 | 0.0000 | None |
| | SV_pbsv_slop25percent | None | 0.0000 | None | None | 0.0000 | None |
| | CNVsandSVs | 0.9003 | 0.9620 | 0.9301 | 0.8686 | 0.8988 | 0.8834 |
| XY | chrX_PAR | 0.0000 | 0.0000 | None | 0.0000 | 0.0000 | None |
| | chrX_XTR | 0.0000 | 0.0000 | None | 0.0000 | 0.0000 | None |
| | chrX_ampliconic | 0.0000 | 0.0000 | None | 0.0000 | 0.0000 | None |
| | chrX_nonPAR | 0.0000 | 0.0000 | None | 0.0000 | 0.0000 | None |
| | chrY_PAR | 0.0000 | 0.0000 | None | 0.0000 | 0.0000 | None |
| | chrY_XTR | 0.0000 | 0.0000 | None | 0.0000 | 0.0000 | None |
| | chrY_ampliconic | 0.0000 | 0.0000 | None | 0.0000 | 0.0000 | None |
| | chrY_nonPAR | 0.0000 | 0.0000 | None | 0.0000 | 0.0000 | None |
| | AllAutosomes | 0.9951 | 0.9928 | 0.9940 | 0.9943 | 0.9890 | 0.9916 |
| Ancestry | ancestry_AFR | 0.9979 | 0.9981 | 0.9980 | 0.9957 | 0.9927 | 0.9942 |
| | ancestry_AMR | 0.9977 | 0.9981 | 0.9979 | 0.9957 | 0.9914 | 0.9935 |
| | ancestry_EAS | 0.9966 | 0.9971 | 0.9969 | 0.9949 | 0.9913 | 0.9931 |
| | ancestry_EUR | 0.9969 | 0.9979 | 0.9974 | 0.9954 | 0.9917 | 0.9935 |
| | ancestry_Neanderthal | 0.9899 | 0.9976 | 0.9937 | 0.9909 | 0.9922 | 0.9916 |
| | ancestry_SAS | 0.9965 | 0.9982 | 0.9973 | 0.9954 | 0.9922 | 0.9938 |

Table S4. The precision, recall, and F1 score of variant calling using 30x multi-platform data (30x ONT + 30x Illumina), stratifying into all defined genomic regions and based on the variant types.

| | | SNPs | | | Indels | | |
|---|---|---|---|---|---|---|---|
| Stratification Type | Stratification Group | Precision | Recall | F1 score | Precision | Recall | F1 score |
| LowComplexity | AHomopol_gt6bp_Imp_gt10bp | 0.9966 | 0.9920 | 0.9943 | 0.9761 | 0.9612 | 0.9686 |
| | ATR201-10kbp | 0.9986 | 0.9961 | 0.9973 | 0.9794 | 0.9566 | 0.9679 |
| | ATR51-200bp | 0.9949 | 0.9923 | 0.9936 | 0.9584 | 0.9293 | 0.9436 |
| | ATR_gt10kbp | 0.9992 | 0.9992 | 0.9992 | 0.9992 | 0.9992 | 0.9992 |
| | ATR_gt100bp | 0.9972 | 0.9945 | 0.9959 | 0.9630 | 0.9286 | 0.9455 |
| | ATR_lt51bp | 0.9965 | 0.9948 | 0.9956 | 0.9814 | 0.9724 | 0.9769 |
| | ATR_and_Homopol | 0.9964 | 0.9929 | 0.9947 | 0.9756 | 0.9605 | 0.9680 |
| | ATR | 0.9963 | 0.9941 | 0.9952 | 0.9744 | 0.9585 | 0.9664 |
| | notin_ATR | 0.9988 | 0.9962 | 0.9975 | 0.9853 | 0.9748 | 0.9800 |
| | SR_diTR_11-50bp | 0.9914 | 0.9870 | 0.9892 | 0.9725 | 0.9599 | 0.9661 |
| | SR_diTR_51-200bp | 0.8974 | 0.8684 | 0.8827 | 0.8719 | 0.8311 | 0.8510 |
| | SR_diTR_gt200bp | 0.0000 | 0.0000 | None | 0.0000 | 0.0000 | None |
| | SR_Homopol_4-6bp | 0.9986 | 0.9957 | 0.9972 | 0.9866 | 0.9713 | 0.9789 |
| | SR_Homopol_7-11bp | 0.9972 | 0.9937 | 0.9954 | 0.9851 | 0.9647 | 0.9748 |
| | SR_Homopol_gt11bp | 0.9884 | 0.9733 | 0.9808 | 0.9670 | 0.9569 | 0.9619 |
| | SR_Homopol_gt20bp | 0.9858 | 0.9268 | 0.9554 | 0.9009 | 0.8858 | 0.8933 |
| | SR_Imp_Homopol_gt10bp | 0.9947 | 0.9875 | 0.9911 | 0.9711 | 0.9579 | 0.9645 |
| | SR_Imp_Homopol_gt20bp | 0.9949 | 0.9790 | 0.9869 | 0.9467 | 0.9191 | 0.9327 |
| | SR_quadTR_20-50bp | 0.9917 | 0.9820 | 0.9868 | 0.9788 | 0.9617 | 0.9702 |
| | SR_quadTR_51-200bp | 0.8750 | 0.8087 | 0.8405 | 0.8909 | 0.8174 | 0.8526 |
| | SR_quadTR_gt200bp | None | 0.0000 | None | None | 0.0000 | None |
| | SR_triTR_15-50bp | 0.9946 | 0.9931 | 0.9938 | 0.9886 | 0.9741 | 0.9813 |
| | SR_triTR_51-200bp | 0.9992 | 0.8889 | 0.9412 | 0.9538 | 0.8983 | 0.9252 |
| | SR_triTR_gt200bp | None | 0.0000 | None | None | 0.0000 | None |

| | | | | | | | |
|---|---|---|---|---|---|---|---|
| | notAHomopol_gt6bp_Imp_gt10bp | 0.9988 | 0.9962 | 0.9975 | 0.9884 | 0.9797 | 0.9841 |
| | notATR_and_Homopol | 0.9988 | 0.9963 | 0.9976 | 0.9966 | 0.9910 | 0.9938 |
| | satellites | 0.9976 | 0.9953 | 0.9965 | 0.9671 | 0.9393 | 0.9530 |
| | not_satellites | 0.9987 | 0.9961 | 0.9974 | 0.9831 | 0.9717 | 0.9773 |
| SegmentalDuplications | ChainSelf | 0.9889 | 0.9731 | 0.9809 | 0.9787 | 0.9611 | 0.9698 |
| | ChainSelf_gt10kb | 0.9718 | 0.9361 | 0.9536 | 0.9661 | 0.9382 | 0.9519 |
| | gt5SegDups_gt10kb | None | 0.0000 | None | None | 0.0000 | None |
| | notChainSelf | 0.9984 | 0.9973 | 0.9982 | 0.9832 | 0.9721 | 0.9776 |
| | notChainSelf_gt10kb | 0.9984 | 0.9972 | 0.9982 | 0.9832 | 0.9720 | 0.9776 |
| | notSegDup | 0.9985 | 0.9979 | 0.9986 | 0.9833 | 0.9723 | 0.9778 |
| | notSegDups_gt10kb | 0.9985 | 0.9979 | 0.9986 | 0.9833 | 0.9723 | 0.9778 |
| | SegDups | 0.9823 | 0.9488 | 0.9653 | 0.9722 | 0.9416 | 0.9566 |
| | SegDups_gt10kb | 0.9795 | 0.9423 | 0.9605 | 0.9688 | 0.9366 | 0.9524 |
| Mappability | nonUnique_l100 | 0.9895 | 0.9584 | 0.9737 | 0.9730 | 0.9132 | 0.9422 |
| | nonUnique_l250 | 0.9386 | 0.7263 | 0.8189 | 0.8866 | 0.6469 | 0.7480 |
| | LowMap | 0.9895 | 0.9584 | 0.9737 | 0.9730 | 0.9132 | 0.9422 |
| | notLowMap | 0.9984 | 0.9984 | 0.9988 | 0.9832 | 0.9729 | 0.9780 |
| OtherDiffcult | KIR | None | 0.0000 | None | None | 0.0000 | None |
| | collapsed_DUP_FP | 0.7854 | 0.9450 | 0.8578 | 0.7967 | 0.8981 | 0.8444 |
| | ppl_CNV_FP | 0.9626 | 0.8993 | 0.9299 | 0.9414 | 0.8612 | 0.8995 |
| | false_DUP_correct | 0.9000 | 0.2264 | 0.3618 | 0.7143 | 0.1724 | 0.2778 |
| | false_DUP_incorrect | 0.9992 | 0.2500 | 0.4000 | None | 0.0000 | None |
| | LD_discord_haplotypes | 0.9986 | 0.9985 | 0.9989 | 0.9806 | 0.9725 | 0.9765 |
| | gnomeAD_InbreedingCoeff | 0.9223 | 0.9298 | 0.9261 | 0.9548 | 0.9302 | 0.9423 |
| | L1H_gt500 | 0.9988 | 0.9226 | 0.9592 | 0.9880 | 0.7746 | 0.8684 |
| | MHC | 0.9974 | 0.9905 | 0.9939 | 0.9547 | 0.9398 | 0.9472 |
| | VDJ | None | 0.0000 | None | None | 0.0000 | None |
| | contigs_lt500kb | None | 0.0000 | None | None | 0.0000 | None |

| | | | | | | | |
|---|---|---|---|---|---|---|---|
| | gaps_slop15kb | None | 0.0000 | None | None | 0.0000 | None |
| | AllOtherDiff | 0.9848 | 0.9651 | 0.9749 | 0.9614 | 0.9345 | 0.9478 |
| FunctionalRegions | notCDS | 0.9987 | 0.9961 | 0.9974 | 0.9831 | 0.9717 | 0.9773 |
| | CDS | 0.9971 | 0.9931 | 0.9951 | 0.9945 | 0.9595 | 0.9767 |
| FunctionalTechnicallyDifficult Regions | BadPromoters | 0.9992 | 0.9992 | 0.9992 | 0.9844 | 0.8592 | 0.9175 |
| | CMRG_duplicationinKMT2C | 0.9992 | 0.9992 | 0.9992 | None | 0.0000 | None |
| | CMRG_falselyduplicatedgenes | 0.9992 | 0.3103 | 0.4737 | None | 0.0000 | None |
| GCContent | gc15 | 0.9976 | 0.9951 | 0.9963 | 0.9789 | 0.9647 | 0.9718 |
| | gc15-20 | 0.9980 | 0.9951 | 0.9966 | 0.9854 | 0.9739 | 0.9796 |
| | gc20-25 | 0.9987 | 0.9967 | 0.9977 | 0.9878 | 0.9799 | 0.9838 |
| | gc25-30 | 0.9988 | 0.9968 | 0.9978 | 0.9836 | 0.9748 | 0.9792 |
| | gc30-55 | 0.9987 | 0.9961 | 0.9974 | 0.9835 | 0.9726 | 0.9780 |
| | gc55-60 | 0.9987 | 0.9961 | 0.9974 | 0.9787 | 0.9640 | 0.9713 |
| | gc60-65 | 0.9987 | 0.9950 | 0.9968 | 0.9769 | 0.9553 | 0.9660 |
| | gc65-70 | 0.9982 | 0.9944 | 0.9963 | 0.9786 | 0.9505 | 0.9644 |
| | gc70-75 | 0.9984 | 0.9959 | 0.9975 | 0.9816 | 0.9486 | 0.9648 |
| | gc75-80 | 0.9983 | 0.9975 | 0.9983 | 0.9829 | 0.9489 | 0.9656 |
| | gc80-85 | 0.9990 | 0.9966 | 0.9978 | 0.9837 | 0.9237 | 0.9527 |
| | gc85 | 0.9947 | 0.9921 | 0.9934 | 0.9803 | 0.9548 | 0.9674 |
| | gclt25orgcgr65 | 0.9986 | 0.9960 | 0.9973 | 0.9860 | 0.9741 | 0.9800 |
| | gclt30orgcgt55 | 0.9987 | 0.9962 | 0.9975 | 0.9824 | 0.9703 | 0.9763 |
| union | AllDiffRegions | 0.9952 | 0.9840 | 0.9896 | 0.9768 | 0.9617 | 0.9692 |
| | notAllDiffRegions | 0.9987 | 0.9989 | 0.9984 | 0.9980 | 0.9947 | 0.9963 |
| | Alllowmapsegdup | 0.9912 | 0.9669 | 0.9789 | 0.9783 | 0.9378 | 0.9576 |
| | notAlllowmapsegdup | 0.9985 | 0.9986 | 0.9989 | 0.9832 | 0.9728 | 0.9780 |
| GenomeSpecific_HG003 | comphetindel10bp | 0.9782 | 0.9818 | 0.9800 | 0.9349 | 0.9081 | 0.9213 |
| | comphetsnp10bp | 0.9971 | 0.9935 | 0.9953 | 0.9396 | 0.9300 | 0.9348 |
| | complexindel10bp | 0.9912 | 0.9841 | 0.9877 | 0.9766 | 0.9379 | 0.9569 |

| | | | | | | | |
|---|---|---|---|---|---|---|---|
| | complexandSVs_AllDiff | 0.9964 | 0.9877 | 0.9920 | 0.9773 | 0.9624 | 0.9698 |
| | complexandSVs | 0.9977 | 0.9875 | 0.9926 | 0.9494 | 0.9179 | 0.9334 |
| | othercomplexwithin10bp | 0.9967 | 0.9637 | 0.9800 | 0.9728 | 0.9141 | 0.9425 |
| | notComplexandSVs_AllDiff | 0.9987 | 0.9990 | 0.9984 | 0.9983 | 0.9954 | 0.9969 |
| | snpswithin10bp | 0.9988 | 0.9952 | 0.9974 | 0.9733 | 0.9511 | 0.9621 |
| | CNV_CCSandONT_elliptical_outlier | 0.9322 | 0.9565 | 0.9442 | 0.9992 | 0.9992 | 0.9992 |
| | CNV_mrcanavarIllumina_CCShighcov_ ONThighcov_intersection | 0.6667 | 0.8182 | 0.7347 | 0.9992 | 0.9992 | 0.9992 |
| | SV_pbsv_slop25percent | None | 0.0000 | None | None | 0.0000 | None |
| | CNVsandSVs | 0.9913 | 0.9805 | 0.9859 | 0.9003 | 0.8804 | 0.8902 |
| XY | chrX_PAR | 0.0000 | 0.0000 | None | 0.0000 | 0.0000 | None |
| | chrX_XTR | 0.0000 | 0.0000 | None | 0.0000 | 0.0000 | None |
| | chrX_ampliconic | 0.0000 | 0.0000 | None | 0.0000 | 0.0000 | None |
| | chrX_nonPAR | 0.0000 | 0.0000 | None | 0.0000 | 0.0000 | None |
| | chrY_PAR | 0.0000 | 0.0000 | None | 0.0000 | 0.0000 | None |
| | chrY_XTR | 0.0000 | 0.0000 | None | 0.0000 | 0.0000 | None |
| | chrY_ampliconic | 0.0000 | 0.0000 | None | 0.0000 | 0.0000 | None |
| | chrY_nonPAR | 0.0000 | 0.0000 | None | 0.0000 | 0.0000 | None |
| | AllAutosomes | 0.9987 | 0.9961 | 0.9974 | 0.9831 | 0.9717 | 0.9773 |
| Ancestry | ancestry_AFR | 0.9985 | 0.9984 | 0.9989 | 0.9847 | 0.9765 | 0.9806 |
| | ancestry_AMR | 0.9983 | 0.9983 | 0.9987 | 0.9840 | 0.9731 | 0.9785 |
| | ancestry_EAS | 0.9983 | 0.9979 | 0.9985 | 0.9832 | 0.9711 | 0.9771 |
| | ancestry_EUR | 0.9984 | 0.9982 | 0.9987 | 0.9840 | 0.9723 | 0.9781 |
| | ancestry_Neanderthal | 0.9991 | 0.9986 | 0.9988 | 0.9845 | 0.9791 | 0.9818 |
| | ancestry_SAS | 0.9987 | 0.9985 | 0.9986 | 0.9847 | 0.9741 | 0.9793 |

### Supplementary Note

#### Input of the pileup network

The input of the pileup network has 18 features, A+, C+, G+, T+, $I_s$+, $I^1_s$+, $D_s$+, $D^1_s$+, $D_R$+, A−, C−, G−, T−, $I_s$−, $I^1_s$−, $D_s$−, $D^1_s$− and $D_R$−, where + and − denote the positive and negative strands, respectively. A, C, G, T, I and D represent the read count supporting the four nucleotides, insertion, and deletion. The subscript 's' means the starting position of an indel, and the subscript 'R' means the following positions of an indel. The superscript '1' means only the Indel with the highest read support is counted when various lengths are observed at a position.

#### Input channels of the full-alignment network

- Reference base: An integer {A:100, G:75, T:50, C:25} is assigned to a position according to the reference base. 0 is used for deleted bases.
- Alternative base: An integer {A:100, G:75, T:50, C:25, insertion starting position: -50, deletion starting position: -100, deletion non-starting positions: 0} is assigned to a position if the alignment at the position mismatches the reference base. 0 is assigned if a position matches the reference base.
- Strand information: An integer {+: 100, -:50} is assigned based on which strand all positions of a read are aligned to. 0 is assigned if the position is in a deletion.
- Mapping quality: An integer ranging from 0 to 100 scaled up from the Phred mapping score (0 to 60, capped at 60 if >60) of an aligned read is assigned to all positions of a whole read. 0 is assigned to deleted bases. Decimal places are truncated.
- Base quality: An integer ranging from 0 to 100 scaled up from the Phred base quality score (0 to 40, capped at 40 if >40) is assigned to each aligned base. 0 is assigned to deleted bases. Decimal places are truncated.
- Candidate proportion: An integer ranging from 0 to 100 to indicate the percentage of reads among all reads that support an exact mismatch pattern in a read is assigned to all positions of the read. 0 is assigned to deleted bases. For example, a candidate with 20 reads, 10 supporting reference alleles A, 5 supporting alternative alleles C, 3 supporting alternative alleles T, and 2 supporting an insertion AT. The integer assigned to the reads that support A, C, T, and AT are 0, 25, 15, and 10, respectively. Decimal places are truncated.
- Insertion base: Inserted bases are exhibited after each insertion's starting position with integers {A:100, G:75, T:50, C:25}. Although it rarely happens, it is possible for exhibited insertions to have overlaps. But an overlap can be disambiguated using the "alternative base" channel, in which the insertion starting positions are assigned -50.
- Phasing information: For unphased reads, the integer 60 is assigned to all reads. For phased reads, {HP1: 30, unphased: 60, HP2: 90} is assigned to all positions of each read. 0 is assigned to deleted bases. If the reads are phased, the reads in all eight channels are sorted in the order "unphased, HP1, HP2".
- (optional) Stratification information: An integer {in the region: 50, not in the region: 100} is assigned based on if the position of variant candidate is in a designated special bed file.

#### Outputs of the pileup and full-alignment network

Both network outputs contain 21 genotype and zygosity tasks, while the full-alignment network contains two more tasks for the length of two indel alleles. The 21-genotype probabilistic model compromises all the possible genotypes of a diploid genome at a position. These genotypes are 'AA', 'AC', 'AG', 'AT', 'CC', 'CG', 'CT', 'GG', 'GT', 'TT', 'AI', 'CI', 'GI', 'TI', 'AD', 'CD', 'GD', 'TD', 'II', 'DD', and 'ID', where ‘A’, ‘C’, ‘G’, ‘T’, ‘I’ (insertion) and ‘D’ (deletion) indicate the six possible alleles. The zygosity task generates the possibility of one of the following: 1) a homozygous reference (0/0), 2) heterozygous with 1 or 2 alternative alleles (0/1 or 1/2), and 3) a homozygous variant (1/1). The indel tasks in the full-alignment network output two indel alleles with their length. Each task outputs the 33 possibilities of the indel length: 1) a longer than 15 bp deletion (<-15 bp), 2) an insertion or deletion with between -15 bp and 15 bp long and 3) a longer than 15 bp insertion (>15 bp).

#### Data Sources

##### Reference genomes

GRCh38_no_alt

https://ftp.ncbi.nlm.nih.gov/genomes/all/GCA/000/001/405/GCA_000001405.15_GRCh38/seqs_for_alignment_pipelines.ucsc_ids/GCA_000001405.15_GRCh38_no_alt_analysis_set.fna.gz

##### GIAB Truth Variants

HG002 (NA24385), GRCh38, v4.2.1

https://ftp-trace.ncbi.nlm.nih.gov/giab/ftp/release/AshkenazimTrio/HG002_NA24385_son/NISTv4.2.1/GRCh38/

HG003 (NA24149), GRCh38, v4.2.1

https://ftp-trace.ncbi.nlm.nih.gov/giab/ftp/release/AshkenazimTrio/HG003_NA24149_father/NISTv4.2.1/GRCh38/

HG004 (NA24143), GRCh38, v4.2.1

https://ftp-trace.ncbi.nlm.nih.gov/giab/ftp/release/AshkenazimTrio/HG004_NA24143_mother/NISTv4.2.1/GRCh38/

##### ONT Sequencing Data

HG002 HD (NA24385), GRCh38_no_alt, ~65x

https://s3-us-west-2.amazonaws.com/human-pangenomics/index.html?prefix=NHGRI_UCSC_panel/HG002/nanopore/

HG003 (NA24149), GRCh38_no_alt, ~77x

https://s3-us-west-2.amazonaws.com/human-pangenomics/index.html?prefix=NHGRI_UCSC_panel/HG003/nanopore/

HG004 (NA24143), GRCh38_no_alt, ~80x

https://s3-us-west-2.amazonaws.com/human-pangenomics/index.html?prefix=NHGRI_UCSC_panel/HG004/nanopore/

##### Illumina Sequencing Data

HG002 PrecisionFDA (NA24385), GRCh38_no_alt, ~35-fold

https://opendata.nist.gov/pdrsrv/mds2-2336/input_fastqs/HG002.novaseq.pcr-free.35x.R1.fastq.gz

https://opendata.nist.gov/pdrsrv/mds2-2336/input_fastqs/HG002.novaseq.pcr-free.35x.R2.fastq.gz

HG003 PrecisionFDA (NA24149), GRCh38_no_alt, ~35-fold

https://opendata.nist.gov/pdrsrv/mds2-2336/input_fastqs/HG003.novaseq.pcr-free.35x.R1.fastq.gz

https://opendata.nist.gov/pdrsrv/mds2-2336/input_fastqs/HG003.novaseq.pcr-free.35x.R2.fastq.gz

HG004 PrecisionFDA (NA24143), GRCh38_no_alt, ~35-fold

https://opendata.nist.gov/pdrsrv/mds2-2336/input_fastqs/HG004.novaseq.pcr-free.35x.R1.fastq.gz

https://opendata.nist.gov/pdrsrv/mds2-2336/input_fastqs/HG004.novaseq.pcr-free.35x.R2.fastq.gz

##### PacBio Sequencing Data

HG002 PrecisionFDA (NA24385), GRCh38_no_alt, ~35-fold

https://nist-midas.s3.amazonaws.com/pdrsrv/mds2-2336/input_fastqs/HG002_35x_PacBio_14kb-15kb.fastq.gz

HG003 PrecisionFDA (NA24149), GRCh38_no_alt, ~35-fold

https://nist-midas.s3.amazonaws.com/pdrsrv/mds2-2336/input_fastqs/HG003_35x_PacBio_14kb-15kb.fastq.gz

HG004 PrecisionFDA (NA24143), GRCh38_no_alt, ~35-old

https://nist-midas.s3.amazonaws.com/pdrsrv/mds2-2336/input_fastqs/HG004_35x_PacBio_14kb-15kb.fastq.gz

#### Commands

##### Read alignment using Minimap2 (v2.17-r941)

```
# Align ONT reads to GRCh38_no_alt

minimap2 -x map-ont -t ${THREADS} -a -A 2 -B 4 -O 4,24 -E 2,1 ref.mmi input.fastq.gz | samtools view -bh -o output.unsorted.bam

samtools sort -@ ${THREADS} -o output.sorted.bam output.unsorted.bam

samtools index -@ ${THREADS} output.sorted.bam
```

##### Read alignment using BWA (v0.7.17-r1198)

```
# Align Illumina reads to GRCh38_no_alt

bwa mem -M -t ${THREADS} ${REFERENCE}.fasta input.R1.fastq.gz input.R2.fastq.gz | samtools view -bh -o output.unsorted.bam

samtools sort -@ ${THREADS} -o output.sorted.bam output.unsorted.bam

samtools index -@ ${THREADS} output.sorted.bam
```

##### Read alignment using pbmm2 (v1.3.0)

```
# Align PacBio reads to GRCh38_no_alt

pbmm2 align --num-threads ${THREADS} --preset CCS --sample ${SAMPLE_NAME} --log-level INFO --sort --unmapped -c 0 -y 70 ${REFERENCE}.fasta input.fastq.gz output.sorted.bam

samtools index -@ ${THREADS} output.sorted.bam
```

#### BAM subsampling using Samtools (v1.10)

### ${FRAC} is the fraction that needs to be subsampled; ${RANDOM_SEED} is the random seed for subsampling

samtools view -@ ${THREADS} -s ${FRAC}.${RANDOM_SEED} -b -o subsampled.bam ${BAM}

samtools index -@ ${THREADS} subsampled.bam

#### Coverage calculation using Mosdepth (v0.2.9)

Mosdepth -t ${THREADS} -n -x –quantize 0:15:150 output ${BAM}

#### Clair3-MP model training

Training section at Clair3-MP github repo.

#### Running Clair3-MP

${CLAIR3-MP_DIR}/run_clair3_mp.sh \

--bam_fn_c=${BAM_PLATFORM_A} \

--bam_fn_p1=${BAM_PLATFORM_B} \

--bam_fn_c_platform=${PLATFORM_A} \

--bam_fn_p1_platform=${PLATFORM_B} \

--output=${OUTPUT_DIR} \

--ref_fn=${REF} \

--threads=${THREADS} \

--model_path_clair3_c=${MODEL_DIR_CLAIR3_PILEUP_FOR_PLATFORM_A} \

--model_path_clair3_p1=${MODEL_DIR_CLAIR3_PILEUP_FOR_PLATFORM_B} \

--trio_model_prefix=${MP_MODEL_PREFIX} \

--model_path_clair3_trio=${MODEL_DIR_CLAIR3_MP_MODEL_FOR_PLATFORM_A_AND_B} \

--sample_name_c=${SAMPLE_PLATFORM_NAME} \

--sample_name_p1=${SAMPLE_PLATFORM_NAME}

#### Running Clair3

docker run -it \

-v ${INPUT_DIR}:${INPUT_DIR} \

-v ${OUTPUT_DIR}:${OUTPUT_DIR} \

-v ${REF_DIR}:${REF_DIR} \

hkubal/clair3:latest \

/opt/bin/run_clair3.sh \

--bam_fn=${INPUT_DIR}/${BAM} \

--ref_fn=${REF} \

```
--threads=${THREADS} \
--platform=${PLATFORM} \
--model_path=/opt/models/${MODEL_NAME} \
--output=${OUTPUT_DIR}
```

##### Benchmarking using hap.py (v0.3.12)

```
hap.py ${GIAB_BASELINE_VCF} output.vcf.gz \
-o ${OUTPUT_DIR}/happy \
-r ${REF} \
-f ${GIAB_CONFIDENT_BED} \
--threads ${THREADS} \
--pass-only \
--engine=vcfeval
```

##### Benchmarking with stratification (v3.1) in hap.py (v0.3.12)

```
qfy.py \
happy.vcf.gz \
-t ga4gh \
-- stratification v3.1-GRCh38-stratifications.tsv \
-o report \
-r ${REF} \
--threads ${THREADS}
```
